# Splam: a deep-learning-based splice site predictor that improves spliced alignments

**DOI:** 10.1101/2023.07.27.550754

**Authors:** Kuan-Hao Chao, Alan Mao, Steven L Salzberg, Mihaela Pertea

## Abstract

The process of splicing messenger RNA to remove introns plays a central role in creating genes and gene variants. Here we describe Splam, a novel method for predicting splice junctions in DNA based on deep residual convolutional neural networks. Unlike some previous models, Splam looks at a relatively limited window of 400 base pairs flanking each splice site, motivated by the observation that the biological process of splicing relies primarily on signals within this window. Additionally, Splam introduces the idea of training the network on donor and acceptor pairs together, based on the principle that the splicing machinery recognizes both ends of each intron at once. We compare Splam’s accuracy to recent state-of-the-art splice site prediction methods, particularly SpliceAI, another method that uses deep neural networks. Our results show that Splam is consistently more accurate than SpliceAI, with an overall accuracy of 96% at predicting human splice junctions. Splam generalizes even to non-human species, including distant ones like the flowering plant *Arabidopsis thaliana*. Finally, we demonstrate the use of Splam on a novel application: processing the spliced alignments of RNA-seq data to identify and eliminate errors. We show that when used in this manner, Splam yields substantial improvements in the accuracy of downstream transcriptome analysis of both poly(A) and ribo-depleted RNA-seq libraries. Overall, Splam offers a faster and more accurate approach to detecting splice junctions, while also providing a reliable and efficient solution for cleaning up erroneous spliced alignments.

## 1. Introduction

Ever since the discovery of RNA splicing ^1^, our understanding of gene transcription and expression has grown steadily more complex. Alternative splicing (AS), where exons and introns are spliced in different combinations to create multiple gene isoforms, is a fundamental mechanism for increasing the diversity of gene function. We now know that the vast majority of human multi-exon genes undergo at least some alternative splicing ^2, 3, 4, 5, 6, 7^.

Recognizing splice sites computationally is a crucial step in accurately assembling gene transcripts and annotating genomes. The most direct method for finding splice sites is to use spliced alignment programs such as TopHat ^8^, SpliceMap ^9^, MapSplice ^10^, STAR ^11^, and HISAT2 ^12^ to align RNA-seq reads directly onto a reference genome. These alignments provide clear evidence of the locations where introns have been spliced out of messenger RNA. However, splice junctions predicted from read alignments are not always reliable, and they may contain false positives due to alignment errors, as well as transcriptional and splicing noise ^13^. Computational tools based on machine learning methods such as GeneSplicer ^14^, MaxEntScan ^15^, SplicePort ^16^, SpliceMachine ^17^, and others ^18^, can help identify some of these false positives. These tools are trained on known splice sites from a target species and are able to recognize splice sites based on the properties of the genome sequence alone. In recent years, convolutional neural networks (CNNs), which have been highly successful in a variety of areas (e.g., ^19, 20, 21, 22^) have been explored as a possible approach to splice site recognition. Several new splice site predictors, including SpliceAI ^23^, SpliceFinder ^24^, Spliceator ^25^, and SpliceRover ^26^, have adopted CNN-based architectures. Attention-based deep learning models have also been recently introduced into the field of splice site prediction, inspired by the success of Google’s Transformer architecture ^27^. Two examples of these models are DNABERT ^28^ and Nucleotide Transformer ^29^. Among these newer methods, SpliceAI is considered to represent the state of the art because it has the highest accuracy ^23^.

In this paper, we introduce Splam, a new deep learning approach that recognizes splice sites from genomic sequences alone more accurately than previous methods. Compared to SpliceAI, our method uses a much shorter flanking context sequence to predict the splice sites, making our model more biologically realistic. In our experiments here, we demonstrate Splam’s accuracy and show how it can be used to improve the accuracy of genome-guided transcript assemblies by removing spurious alignments produced by spliced aligners.

## 2. Results

### 2.1 Splam: an accurate splice junction recognition model

We implemented Splam using a framework based on deep convolutional neural networks to predict splice junctions. The design is illustrated in Figure 1a and is discussed in more detail in <u>Methods</u>. As illustrated in Figure 1b and c, the input to Splam is a DNA sequence composed of 200 nucleotides (nt) on both sides of the donor and acceptor sites, totaling 800nt, and the output is the probability for every base pair of being a donor site, an acceptor site, or neither (see <u>Methods</u>).

**Figure 1:**
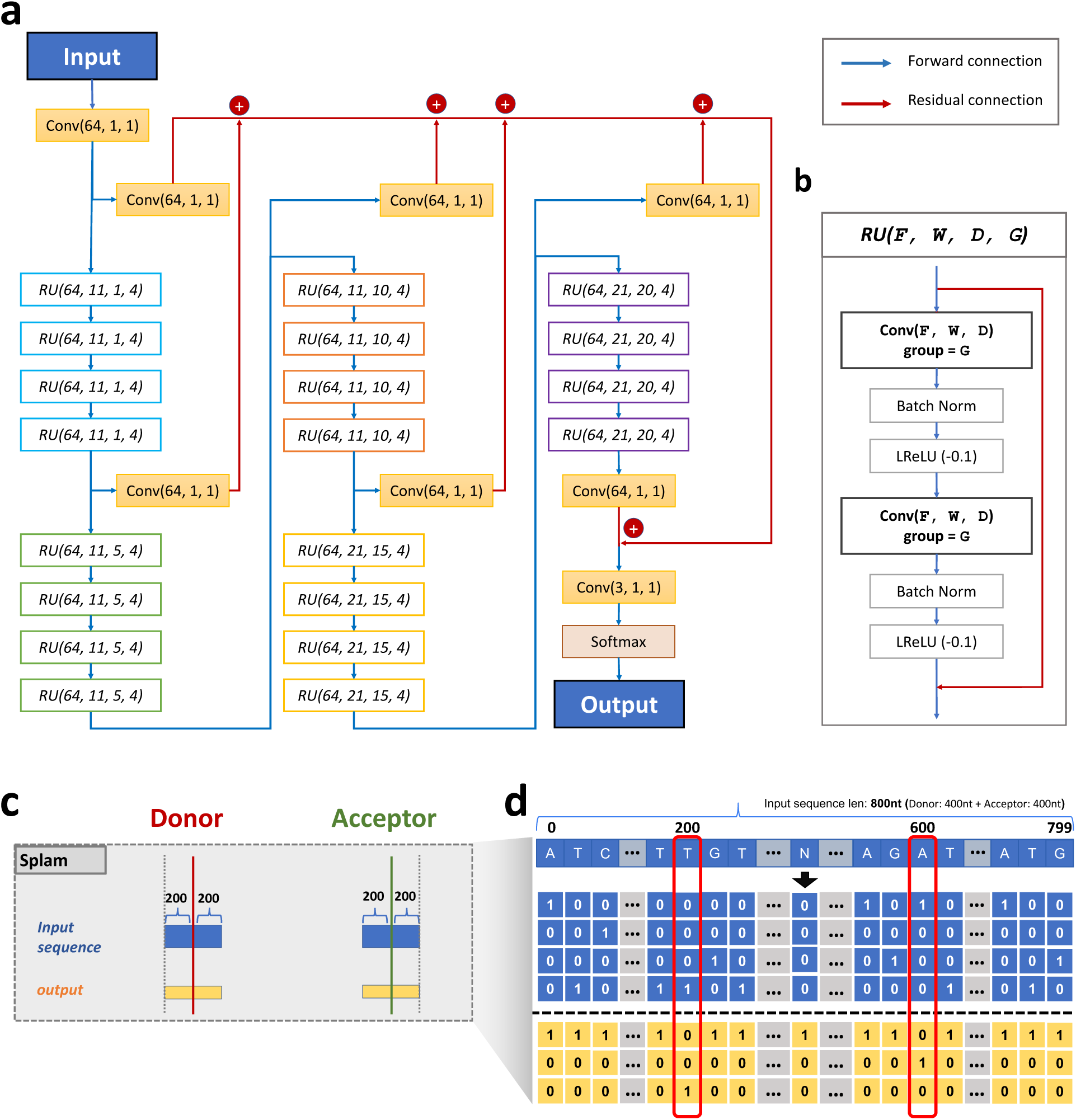
**(a)** The Splam model architecture consists of 20 residual units (RU), where **(b)** each RU comprises two convolutional layers. The convolutional layers utilize a parameter group of 4, along with batch normalization and leaky rectified linear unit (LReLU) activations. Blue arrows indicate forward connections, while red arrows represent residual connections. Within each RU, the input is connected to the output through a residual connection. Overall, the Splam model contains a total of 651,715 parameters. **(c)** Splam’s inputs and outputs. The input sequence (in blue) uses 400nt flanking the donor site and another 400nt flanking the acceptor site. The output (in yellow) is an array of labels for all 800nt. **(d)** The one-hot encoding procedure for the inputs and outputs of Splam employed during model training and testing with *A* represented as [1,0,0,0], *C* as [0,1,0,0], *G* as [0,0,1,0], *T* as [0,0,0,1], and *N* as [0,0,0,0]. Each pair of donor and acceptor sites is concatenated into an 800nt sequence as shown, and the 200^th^ and 600^th^ nucleotides are labeled as [0, 1, 0] (for a donor site) and [0, 0, 1] (for an acceptor site) respectively.

The Splam algorithm focuses on training the model to recognize splice junction patterns at the “splice junction” level; i.e., it attempts to recognize donor and acceptor sites in pairs, just as the spliceosome operates in the cell when it splices out an intron. Note that most human genes have multiple splice isoforms, with current annotation catalogs containing 5-10 isoforms per gene. Thus, an accurate model of splicing should be able to predict all of the introns in these multiple variants. Splam demonstrates this ability, as our experiments will show.

In contrast, the current state-of-the-art tool, SpliceAI, was trained on a single isoform per gene, and moreover was trained on protein-coding genes only. It was originally tested by measuring its ability to predict a single set of splice sites from the canonical isoform for each protein-coding gene. Designing a splice site recognition method based only on one transcript per gene may result in mislabeling alternative splice sites even when they are perfectly valid. Our experiments sought to test the hypothesis that our training regimen, which uses the splice sites from multiple isoforms for each gene, might result in better predictions.

To assess the accuracy of Splam versus SpliceAI in recognizing splice junctions, we created two distinct inputs to SpliceAI for our experiments. The first, simply called SpliceAI, provided 10,400nt of sequence for each splicing event, following the original SpliceAI study’s approach, which included 200nt upstream and downstream of the donor and acceptor sites plus 5,000nt of flanking sequence on both sides of each sequence being scored. For the second set of inputs, called SpliceAI-10k-Ns, only 400nt of flanking sequence was provided (along with the entire sequence of the intron). The precise inputs are shown in <u>Figure S1a</u> and discussed in <u>Methods</u>. <u>Figure S1b</u> shows the relative amount of sequence data used for prediction by Splam, SpliceAI, and SpliceAI-10k-Ns. As the figure illustrates, SpliceAI uses far more context to predict splice sites, even when we replace its normal 10 kilobase (Kb) flanking input with Ns, because it uses the entire length of the intron. For our experiments, SpliceAI had 14–20 times as many base pairs available to make predictions, but despite this advantage, Splam had superior accuracy, as we demonstrate below.

In all experiments described here, programs were trained on genes from all chromosomes except Chr1 and Chr9, and then tested on genes from chromosomes 1 and 9, after first removing homologous sequences (see <u>Methods</u>). This mirrors the training strategy for SpliceAI, which also trained on genes from all chromosomes except Chr1 and Chr9 and then tested on chromosomes 1 and 9.

We computed the accuracy of distinguishing positive splice junctions in randomly selected subsets of two datasets, designated `Positive-MANE` and `Positive-Alt`, from a random subset of negative examples of splice junctions in the `Negative-1` and `Negative-Random` datasets. `Positive-MANE` comprises splice junctions selected from the MANE annotation ^30^ that are supported by a minimum of 100 alignments in a large collection of RNA-seq samples from GTEx. `Positive-Alt` consists of splice junctions present in protein-coding genes from the RefSeq ^31^ annotation but not in MANE, and also supported by at least 100 alignments. Thus, all positive examples derive from protein-coding genes, and for many genes, the data includes more than one isoform. `Negative-1` contains splice junctions on the opposite strand of annotated gene loci, supported by exactly 1 alignment, while `Negative-Random` comprises random GT-AG pairs on the opposite strand of annotated genes (see <u>Methods</u> for more details).

The range of precision-recall values at different thresholds obtained by Splam and SpliceAI are shown in Figure 2. Figure 2a displays receiver operating characteristic curves (ROC) and precision-recall curves (PRC) for both Splam and SpliceAI at the junction level. It demonstrates that Splam and SpliceAI perform nearly identically on splice sites from the MANE database but show a noticeable difference in predicting alternative splice sites, with Splam performing better. Furthermore, when SpliceAI is not provided the 10Kb flanking sequence, its performance drops noticeably. Thus, in this data, Splam predicts splice junctions in non-canonical transcripts (`Positive-Alt`) with far fewer false negatives than SpliceAI and SpliceAI-10k-Ns.

**Figure 2:**
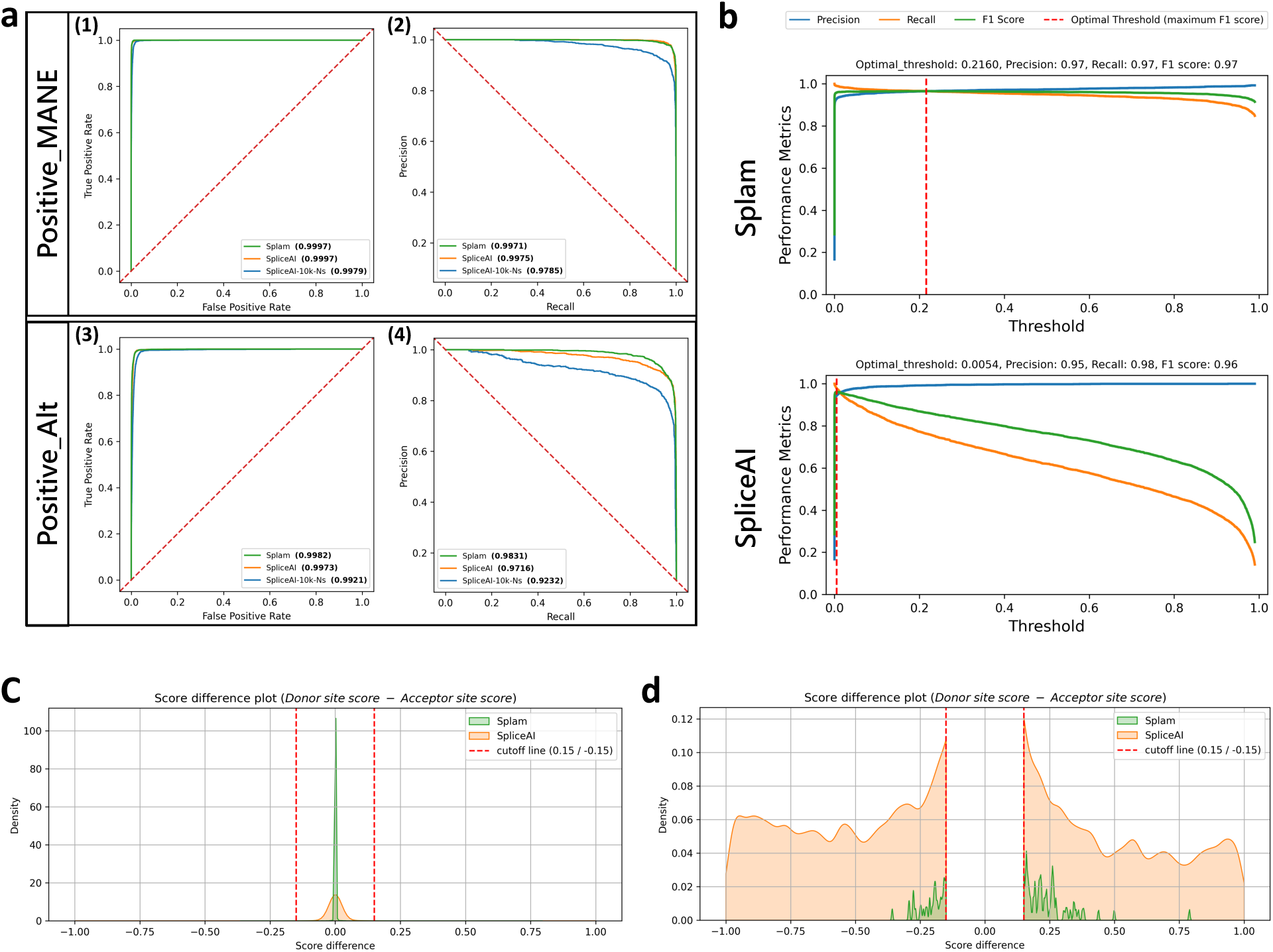
**(a)** ROC and PR curves on two test sets of splice junctions from human chromosomes 1 and 9. Results are shown at the junction level, where the junction score is determined by the minimum of its donor and acceptor scores. Plots **a1-2** show results when classifying `Positive-MANE` where the negative examples included both `Negative-1` and `Negative-Random`. The test set, Test22K-MANE, consisted of 2,000, 10,000, and 10,000 examples from `Positive-MANE`, `Negative-1`, and `Negative-Random` respectively, yielding a positive-to-negative ratio of one to ten (see Methods). Plots **a3-4** show the results of classifying the test set, Test22K-Alt, consisting of 2,000 `Positive-Alt` examples (splice sites in RefSeq but not in MANE) and the same 20,000 negative examples (see Methods). The green curve represents Splam, the orange curve represents SpliceAI with a 10Kb flanking sequence (SpliceAI), and the blue curve represents SpliceAI where the 10Kb flanking sequence is replaced with Ns (SpliceAI-10k-Ns). AUC-ROC values for **(1)** and **(3)** and AUC-PR values for **(2)** and **(4)** are shown next to each tool’s name. **(b)** Discrimination threshold (DT) plots for Splam and SpliceAI on a test set of 2,000 `Positive-MANE`, 2,000 `Positive-Alt`, 10,000 `Negative-1`, and 10,000 `Negative-Random` examples, using all positive and negative examples from **(a)** (Test24K, see Methods). Each plot shows the precision (blue curve), recall (orange curve), and F1 score (green curve) calculated at different thresholds. The threshold at which the F1 score is maximized is indicated by a red dashed line. **(c-d)** Kernel density plots visualizing the differences between donor and acceptor scores (donor score − acceptor score). In **(d)** the score differences between -0.15 and 0.15 have been removed to provide a closer look at the splice junctions where the scores of donor and acceptor were relatively large. The green density plot represents Splam, and the orange density plot represents SpliceAI.

Although the ROC and PR curves for SpliceAI appear excellent, we observed that the raw scores for true donor and acceptor sites were often very low, even if they were slightly higher than the scores for false sites. Ideally, real splice sites will score close to 1, while false ones will be closer to 0. Figure 2b shows that the optimal threshold for SpliceAI is very small (0.0054) and performance quickly drops as the threshold increases. In contrast, the performance of Splam remains stable under a much wider range of thresholds. For instance, when using a reasonably high score threshold of 0.8, Splam achieved an overall accuracy of 96.1% on a test set of 40,000 randomly selected examples, with equal numbers of true and false splice junctions, called Test40K (see <u>Methods</u>); here, accuracy reflects the proportion of correct predictions (positive and negative) out of all predictions. A junction was considered correct if both the donor and acceptor sites were predicted correctly. Precision, defined as the proportion of predicted positive junctions out of all predictions, was even higher, at 99.6%. Recall, defined as the proportion of true junctions that were correctly predicted, was slightly lower but still high, at 92.5%. The precision and recall rates for donor and acceptor sites considered separately were very similar, at 99.6% and 93.0% for both types of sites. By comparison, at the same threshold, SpliceAI had an overall accuracy of 74.1% for splice junctions, and 82.1%/84.5% for donors and acceptors on the same data (Table 1).

**Table 1:**
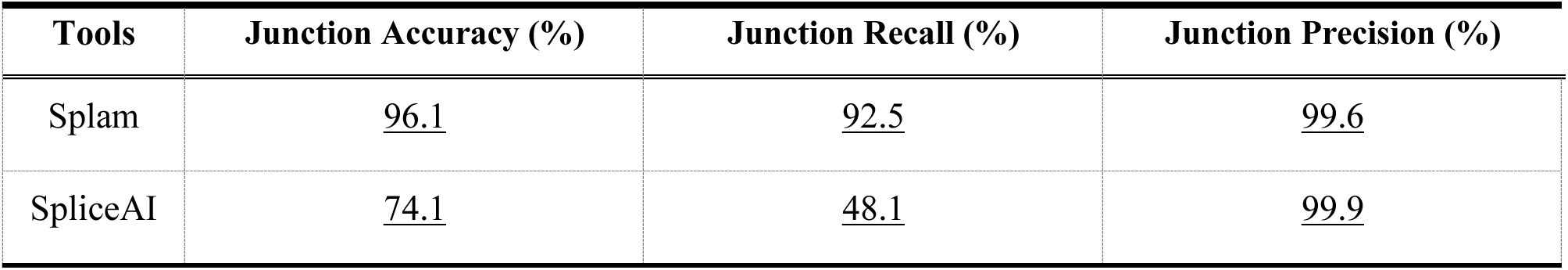
Accuracy, recall, and precision for predicting human splice junctions using a score threshold of 0.8 for Splam and SpliceAI in the Test40K dataset, which contained 20,000 positive examples and 20,000 negative examples (see <u>Methods</u>).

We also investigated whether the donor and acceptor scores were in accord with one another; i.e., if the two scores on both ends of an intron were similar. Figure 2c-d displays the score differences between donor and acceptor sites for Splam and SpliceAI. The plot reveals that although both programs tend to assign very similar scores to both the donor and acceptor flanking an intron, Splam’s scores are in closer agreement. Figure 2d shows a zoomed-in view that illustrates how, in a small fraction of cases, SpliceAI has a score near zero for one site while the other site scores near 1, while the scores for Splam almost never diverge in this manner.

To better understand the score distribution, we investigated the scores produced by Splam and SpliceAI on the Test40K dataset (plotted in <u>Figure S2)</u>. Figure 3 shows the distributions of scores that Splam and SpliceAI assigned to both true and false donor and acceptor sites at a threshold of 0.1. For clarity in the figure, we omitted the points where Splam and SpliceAI agree and are correct, which totaled 16,820 positives and 19,809 negatives, so the figure only shows junctions where at least one program was wrong. The largest class of disagreements were the 2,609 true splice junctions where Splam was correct and SpliceAI was wrong, shown in red. In contrast, there were only 202 true junctions (light blue) where SpliceAI was correct and Splam was wrong. Figure 3a.1 illustrates how the Splam scores cluster into two groups, near (0, 0) and (1, 1), corresponding to splice sites where both donor and acceptor are either low-scoring or high-scoring. As shown in Figure 3a.2, SpliceAI is more likely to assign high scores to one splice site and very low scores to the other; i.e., the junction scores are inconsistent.

**Figure 3:**
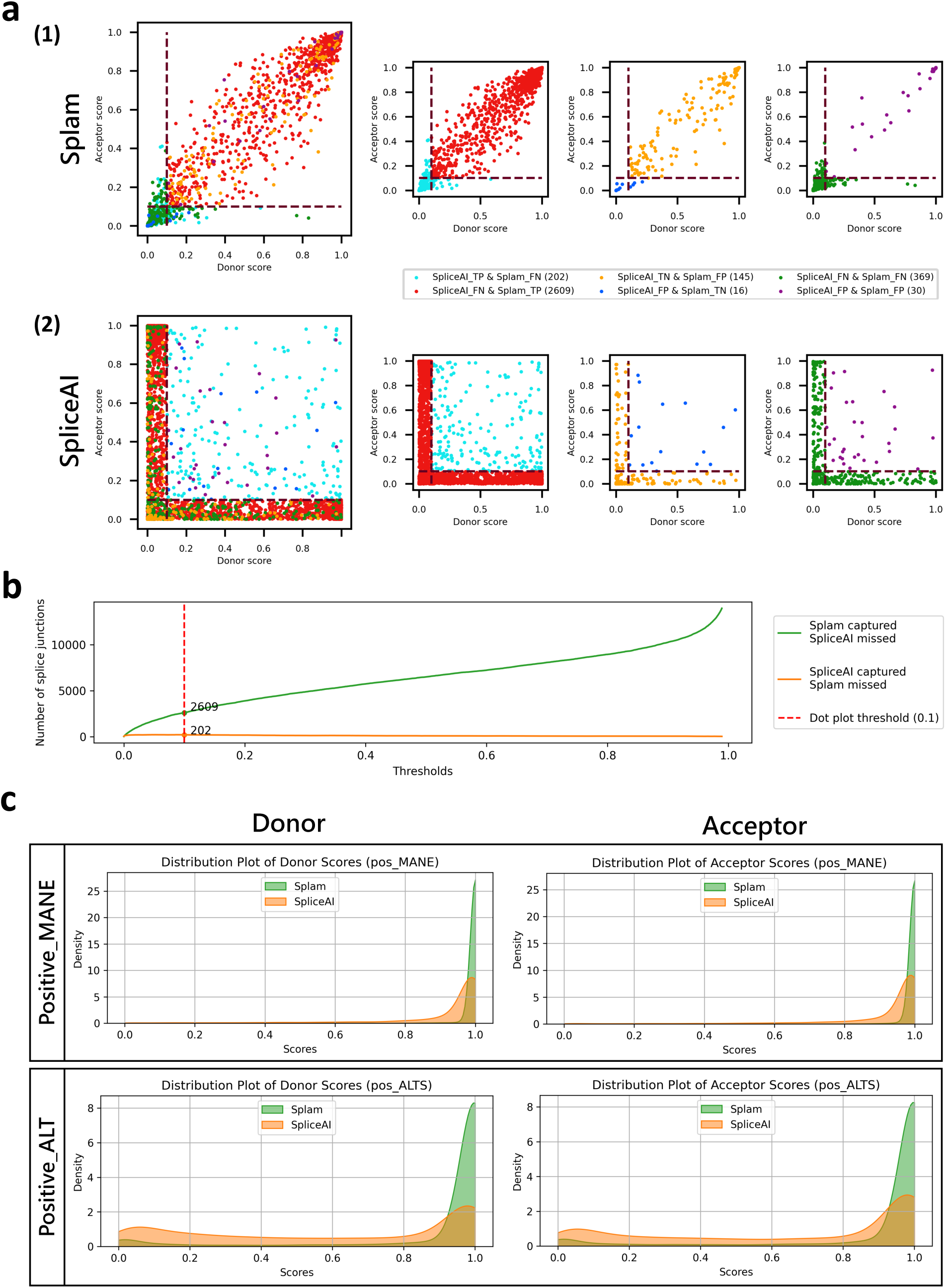
**(a)** Scatter plots for Splam **(1)** and SpliceAI **(2)** on a test set, Test40K, consisting of 10,000 `Positive-MANE`, 10,000 `Positive-Alt`, 10,000 `Negative-1`, and 10,000 `Negative-Random` examples, with each dot representing a splice junction. The red dashed lines show the 0.1 cutoff threshold used here to label splice sites as true or not. The smaller plots in the second and the third columns show subsets of junctions where one program was correct while the other was incorrect (TP and FN, or FP and TN), and the fourth column shows subsets of junctions where both programs made the wrong prediction (either FP and FN). **(b)** Number of splice junctions at all thresholds where Splam and SpliceAI disagree with each other. The green curve represents the instances where Splam correctly predicts them as true positives but SpliceAI predicts them as false negatives; the orange curve represents the instances where SpliceAI correctly predicts them as true positives but Splam predicts them as false negatives. The vertical dashed red line represents the 0.1 threshold used in **(a)**. **(c)** Score density plots for Splam (in green) and SpliceAI (in orange) for donor and acceptor scores on sites found in the MANE dataset (`Positive-MANE`, first row) and sites found in RefSeq but not MANE (`Positive-Alt`, second row).

Although SpliceAI has many more false negatives (missed splice sites), Splam does have slightly more false positives, 145 versus just 16 for SpliceAI. In general, for splice site recognition, we consider the false positives less problematic because the splice-site-like unannotated sequences (i.e., sequences that either program labels as splice sites although they are not annotated as such) are not necessarily true negatives, since they might be recognized and spliced at low levels ^13^. In contrast, failing to identify legitimate positive splice junctions in MANE or RefSeq (false negatives) is much more likely to be a genuine error.

We further focused on positively-labeled splice junctions which were correctly captured by one tool but not the other (Figure 3b). Our findings indicate that Splam consistently captures more splice junctions compared to SpliceAI across all thresholds. Furthermore, Splam outperforms SpliceAI in distinguishing splice junctions in alternatively spliced isoforms, while both programs demonstrate good accuracy in discriminating canonical splice junctions in MANE (refer to Figure 3c-d). Not surprisingly, Splam exhibits even more significant improvements when compared to SpliceAI-10k-Ns, as illustrated in <u>Figure</u> <u>S3</u>, <u>Figure S4</u>, <u>Figure S5</u>, and <u>Figure S6</u>.

### 2.2 Generalization to other species

A frequent concern about deep learning methods is whether they simply memorize the training data, or if their predictive models will work on data that diverges from what they have seen in training. To evaluate the generalization ability of Splam’s model of splicing, we collected data from three successively more distant species and applied Splam to each of them, without re-training. For comparison, we also applied SpliceAI to the same data.

For this experiment, we chose genes from three highly-curated reference genomes: chimpanzee (*Pan troglodytes*), mouse (*Mus musculus*), and the flowering plant *Arabidopsis thaliana*. For each genome, we extracted a random sample of 25,000 introns along with flanking sequences, as defined by the reference annotation (see <u>Methods</u>), and applied both Splam and SpliceAI. The results are shown in Figure 4.

**Figure 4:**
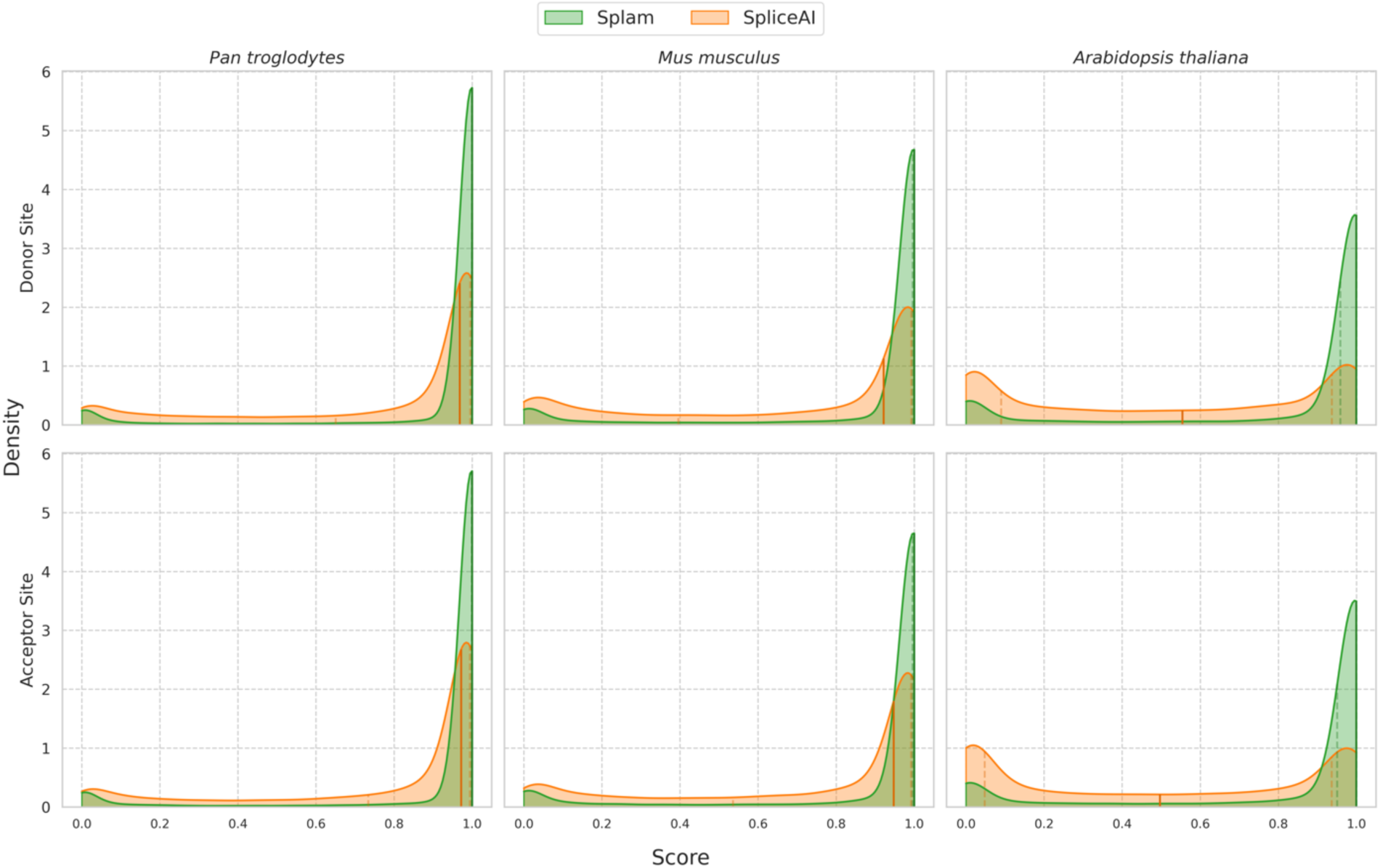
Comparison of score distributions for Splam (green) and SpliceAI (orange) when applied to 25,000 randomly chosen splice sites from chimpanzee (left), mouse (center), and *Arabidopsis* (right). Results for donor sites are shown across the top, and acceptor sites on the bottom. Scores assigned by each program are plotted along the x-axis, while densities for both donor and acceptor sites are plotted on the y-axis. The darkened vertical line through the distribution marks the median value, and the two neighboring dotted lines represent the first and third quartiles. The scores for Splam exhibit a narrow distribution that peaks near 1.0, illustrating that the vast majority of annotated splice sites are assigned high scores, even in the *Arabidopsis* genome. Scores for SpliceAI are more spread out for the chimpanzee and mouse, and the scores for *Arabidopsis* splice sites show an M-shaped distribution peaking at 0 and 1 with a median near 0.5, rather than just peaking near 1.

For both Splam and SpliceAI, the mammalian datasets (chimpanzee and mouse) show similar score distributions to their scores on human splice sites, indicating that both programs achieved some generalization. However, in both cases Splam’s scores show a tighter distribution around 1.0, indicating that it recognizes the non-human splice sites somewhat more strongly. For the plant genome, though, which is far more distant from human, Splam still assigns very high scores to a large majority of splice sites, as shown in the figure, while SpliceAI seems to recognize very few annotated splice sites. Thus despite (or possibly because of) its much smaller input window, Splam’s performance generalizes quite well, even to species as distant as plants.

In addition, we evaluated Splam and SpliceAI using datasets of negative examples for these species (see <u>Methods</u>), and both programs exhibited prominent peaks near 0 (<u>Figure S7</u>). Notably, even in the plant genome, Splam consistently demonstrated low false negatives across different thresholds, outperforming SpliceAI (see ROC and PR curves in <u>Figure S8</u>).

At a score threshold of 0.8, the recall of Splam for splice junctions is 91%, 87%, and 80% for chimpanzee, mouse, and *Arabidopsis* respectively. At the same threshold, SpliceAI had recall rates of 57%, 47%, and 20% (Table 2). The results when using a score threshold of 0.1 are shown in <u>Table S1</u>.

**Table 2:** Accuracy, recall, and precision for donor sites, acceptor sites, and splice junctions at the score threshold of 0.8 for Splam and SpliceAI in chimpanzee *(Pan troglodytes*), mouse *(Mus musculus),* and the flowering plant *Arabidopsis thaliana*.

| Tool |  | Accuracy (%)<br>(Donor / Acceptor / Junction) | Recall (%)<br>(Donor / Acceptor / Junction) | Precision (%)<br>(Donor / Acceptor / Junction) |
| --- | --- | --- | --- | --- |
| *Pan troglodytes* | Splam | 95.7 / 95.7 / 95.5 | 91.4 / 91.4 / 91.0 | 99.9 / 99.9 / 99.9 |
|  | SpliceAI | 84.6 / 86.0 / 78.7 | 69.2 / 72.0 / 57.3 | 99.9 / 99.9 / 99.9 |
| *Mus musculus* | Splam | 93.6 / 93.7 / 93.3 | 87.3 / 87.4 / 86.6 | 99.9 / 99.9 / 99.9 |
|  | SpliceAI | 80.0 / 82.1 / 73.6 | 59.7 / 64.2 / 47.3 | 99.9 / 99.9 / 99.9 |
| *Arabidopsis thaliana* | Splam | 90.7 / 90.4 / 90.0 | 81.4 / 80.9 / 80.2 | 99.9 / 99.9 / 99.9 |
|  | SpliceAI | 68.2 / 67.8 / 60.0 | 36.3 / 35.7 / 20.0 | 99.8 / 99.9 / 99.9 |

Worth noting is that *A. thaliana* has far shorter introns, with an average length of 168nt compared to 6200nt for humans and other mammals. No measurable correlation between intron length and score was observed for either Splam or SpliceAI (<u>Figure S9</u>).

### 2.3 Effect of filtering low-scoring spliced alignments on transcriptome assembly

One crucial step in many transcriptome analyses involves aligning spliced reads to the genome, using an alignment program such as HISAT2 ^12^ or STAR ^11^. The output from these programs might contain two types of erroneous alignments: (1) incorrect alignments due to sequencing errors, repeats, or other sequence artifacts (computational noise), and (2) correct alignments of reads that result from noisy splicing (biological noise). Either computational or biological noise ^32^ can lead to systematic underestimates of transcript abundance levels and large increases in the number of false positive genes and transcripts ^13^. We reasoned that both types of noisy junctions should get low scores from Splam; thus we wanted to evaluate if we could use Splam to improve the quality of transcriptome assemblies by removing low-scoring splice junctions. Using the workflow illustrated in Figure 5 and further detailed in <u>Methods</u>, we processed 20 randomly selected RNA-seq samples, of which 10 were prepared using a poly-A capture library preparation, and 10 were prepared using ribosomal RNA depletion.

**Figure 5:**
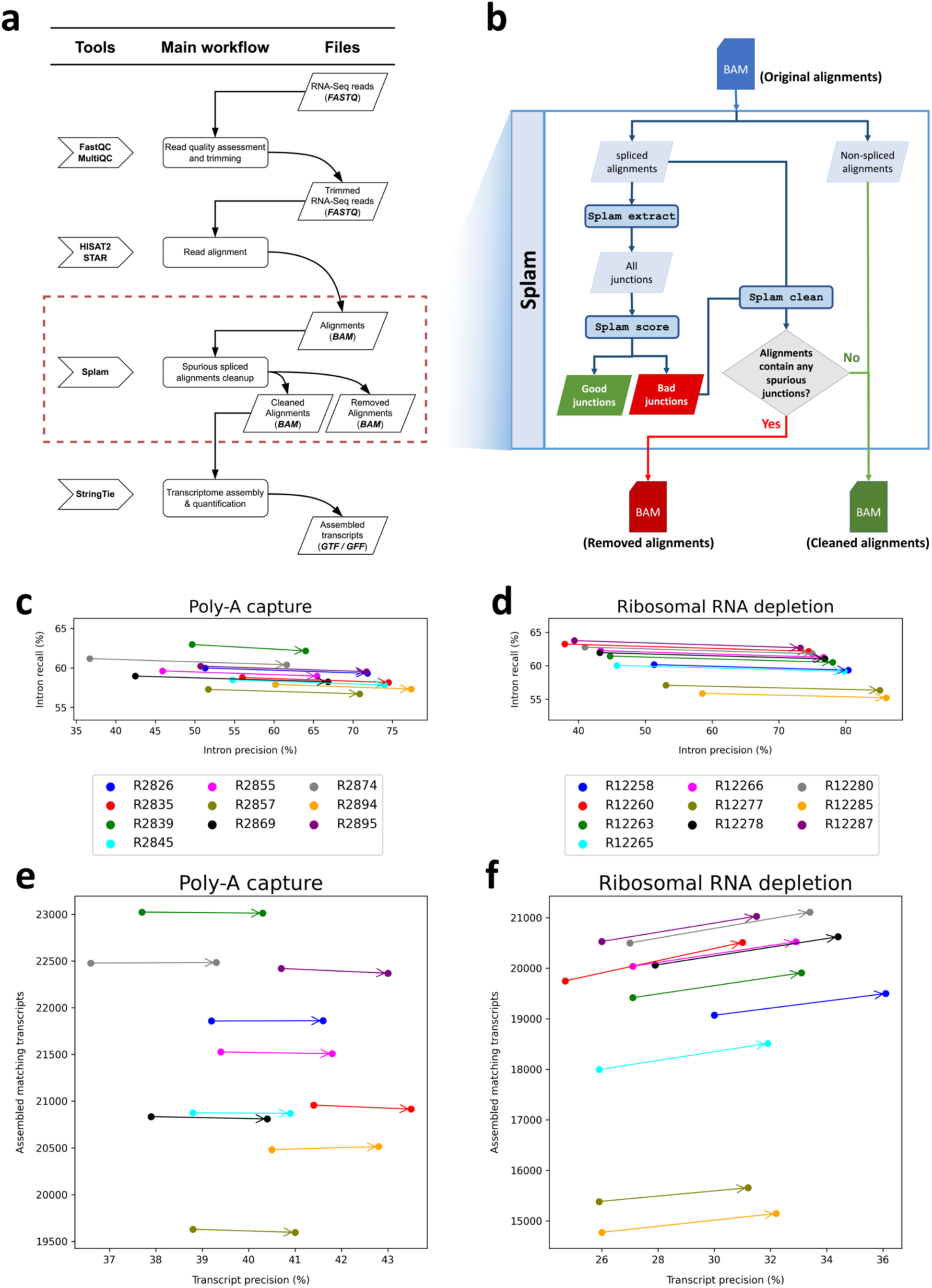
**(a)** The workflow of a typical RNA-Seq pipeline. **(b)** The modified workflow using Splam to remove spurious alignments. **(c)-(f)** Effects of using Splam to remove low-scoring splice junctions from RNA-seq experiments. **(c)-(d)** Results of comparing introns in the RefSeq annotation with splice junctions extracted from the alignment files of 10 poly-A capture **(c)** and 10 rRNA depletion **(d)** RNA-seq samples. Each dot corresponds to a sample, with the original and Splam-cleaned samples connected by an arrow. **(e)-(f)** Results of comparing RefSeq transcripts with assembled transcripts from the same alignment files of poly-A capture **(e)** and rRNA depletion **(f)**, before and after Splam clean-up. As in **(c)** and **(d)**, each dot corresponds to a sample, and the original and Splam-cleaned samples are connected by arrows. The x-axis in all plots shows precision, defined here as the percentage of predicted introns/transcripts that match RefSeq. The y-axis in a-b shows intron recall, defined as the percentage of RefSeq introns that were captured in the files. In **(c)-(d)**, the y-axis shows the number of assembled transcripts that matched RefSeq transcripts.

First, we compared all the splice junctions in the alignment files before and after filtering with Splam to all introns in the RefSeq GRCh38.p14 version 40 annotation file. While some of the splice junctions in the alignment files are no doubt real, an accurate filtering procedure should increase the intron precision, while minimizing the decline in intron recall as computed with respect to RefSeq (see <u>Methods</u>). Indeed, Splam’s results do follow this pattern, as illustrated in Figure 5. Figure 5c and <u>Table S2</u> show the metrics for the 10 poly-A RNA-seq samples. The precision scores for each sample showed a substantial absolute percentage increase, ranging from 14% to 25%, with only a minimal reduction in recall (less than 1%). Figure 5d and <u>Table S3</u> illustrate the results for the 10 rRNA depletion samples, where precision exhibited even higher absolute percentage improvements, ranging from 28% to 37%, with only a slight decrease in recall, ranging from 0.7% to 1.0%.

Second, we evaluated the impact of the Splam removal procedure on downstream transcriptome assembly. We ran StringTie ^33^ on both the original alignment files and the Splam-cleaned alignment files and compared how closely the transcript assemblies match the RefSeq gene annotation file. A successful removal procedure would improve both the precision and recall of transcriptome assembly. Among the 10 poly-A capture samples, the number of matching transcripts remained almost unchanged, while the precision scores consistently improved by an absolute percentage of 2.1% to 2.7% (Figure 5e and <u>Table</u> <u>S4</u>). For the 10 ribosomal RNA depletion samples, both metrics showed even more substantial improvements. Following the Splam removal process, we consistently observed an increase of 500 to 600 matching transcripts being assembled, while precision showed a 5.5% to 6.5% absolute percentage improvement (Figure 5f and <u>Table S5</u>). Other transcriptome assembly metrics, such as the number of matching intron-chains, assembled hypothetical exons, and missed exons from the annotation, show the same picture: by eliminating low-scoring spliced alignments, we achieve more accurate assemblies of transcripts from both poly-A capture and ribosomal RNA depletion RNA-Seq samples, with a particularly notable impact in the latter samples (see <u>Tables S6-S11</u>).

## 3. Discussion

In this study, we presented Splam, a convolutional neural network that recognizes splice junctions in their genomic context. An important principle that we used in designing Splam was that the input should be constrained to be biologically realistic. The previous state-of-the-art CNN-based system, SpliceAI, relies on a window of 10,000 base pairs flanking each splice site plus the entire intronic sequence, which is far larger than what the splicing machinery in the cell can possibly recognize. Introns are flanked by a 5’ donor site and a 3’ acceptor site, usually beginning and ending with the dinucleotides GT and AG. A short window of sequence around each end is recognized by the spliceosome, as well as a branch point about 35-40 bases upstream of the acceptor site ^34^. In addition to these three limited regions, short motifs within the introns and exons known as splicing enhancers and repressors might also affect the splicing process. In total, then, at most a few hundred nucleotides determine whether or not an intron is recognized and spliced out. Notably, all of the crucial sequences involved in splicing and spliceosome assembly are localized in close proximity to the donor and acceptor sites. Thus, it should only be necessary to look in these regions to predict splice sites. We were concerned that the performance improvements observed in SpliceAI when increasing the flanking sequences from 80nt to 10,000nt, as described in ^23^, may have been due to overfitting or memorization of the training set.

In the biological context, donor and acceptor sites are not independent. Therefore, instead of training separately on these two types of splice sites, Splam is specifically designed and trained to recognize pairs of donor and acceptor sites, mimicking the behavior of the spliceosome. Our experiments demonstrate that the biologically-inspired design of Splam produced a highly accurate method, with an overall accuracy of 96% at predicting human splice junctions, exceeding the accuracies of previous methods.

We also showed that the Splam model generalizes quite well, as illustrated by our evaluations on chimpanzee, mouse, and *Arabidopsis*. Unsurprisingly, its performance on chimpanzee was very similar to its performance on human, but perhaps more surprising was its accuracy in predicting splice sites in a plant genome, where it obtained 81.4% sensitivity on donors and 80.9% sensitivity on acceptors. We would expect that if re-trained on species-specific data, Splam would improve even further on those species.

We also tested Splam’s ability to remove spurious splice junctions from the output of a spliced aligner. We showed that when used in this manner, Splam can create alignment files that contain many fewer noisy splicing events while losing only a small number of true splice sites. When the cleaned files are used as input to a transcriptome assembler such as StringTie2, we not only observed higher precision but also an increased number of assembled transcripts that matched known transcripts. These improvements were particularly notable when the filtering process was applied to RNA-seq data generated using rRNA depletion, which generally contains many more incompletely processed transcripts.

## 4. Online Methods

### 4.1 Generating high-quality datasets for splice junctions

We curated two sets of positive examples, “`Positive-MANE`” and “`Positive-Alt`”, and two sets of negative examples, “`Negative-1`” and “`Negative-Random`”, to serve as high-quality datasets for Splam model training and testing, as shown in Figure 6a. The MANE data was taken from the Ensembl/GENCODE GRCh38 MANE release 1.0 file. Note that Refseq and GENCODE agree 100% on all MANE annotations ^30^. To create the `Positive-Alt` dataset, we used RefSeq annotation version 110 ^31^, from which we collected all splice junctions missing from MANE. The resulting annotation file contains information about (1) the location of all annotated splice junctions and (2) the label information of the indices of the donor and acceptor site for each splice junction. We used GRCh38 assembly version 40 as the reference genome sequence.

**Figure 6:**
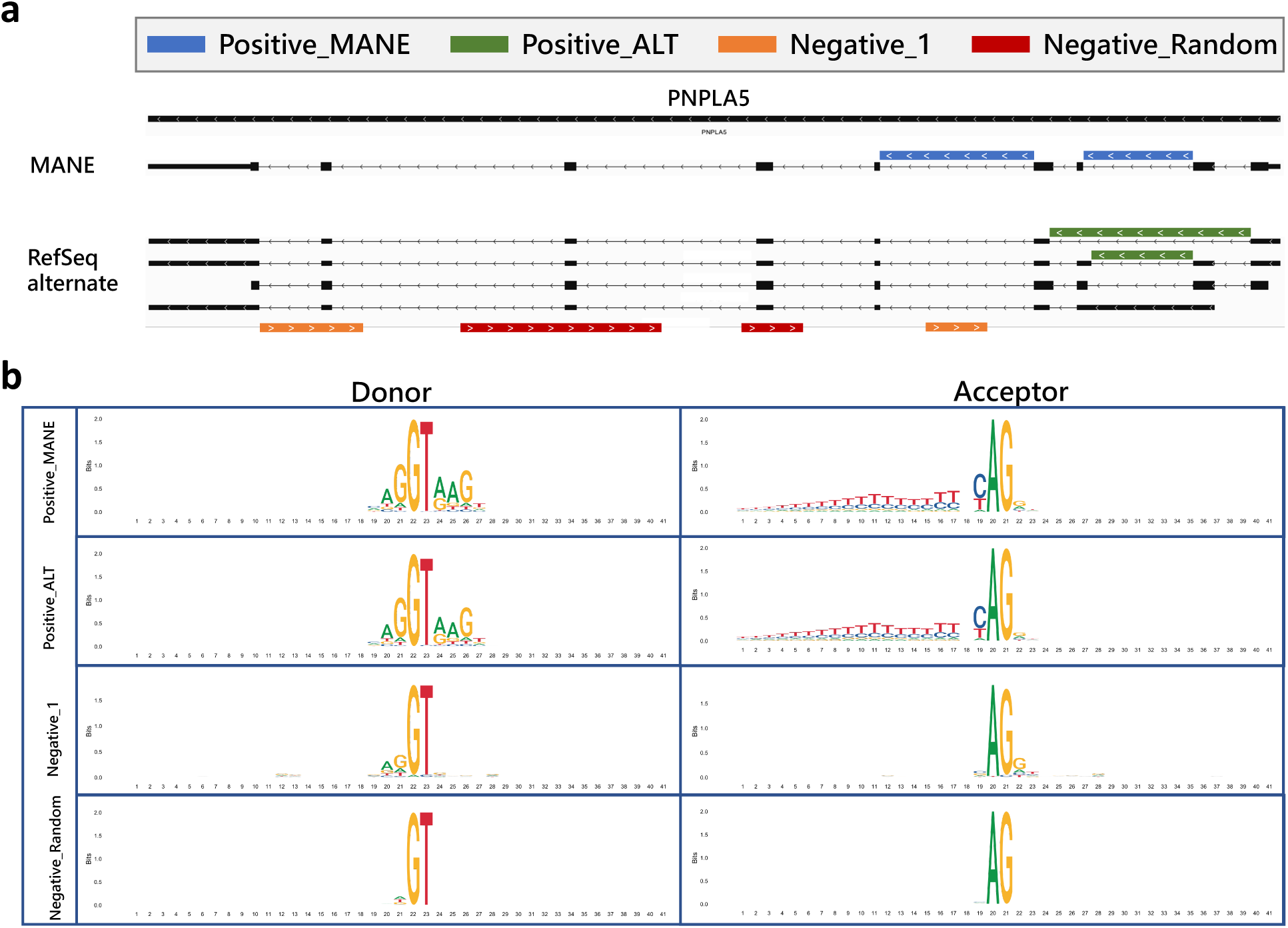
**(a)** Illustration of the four types of splice sites used for training and testing Splam: `Positive-MANE`, `Positive-Alt`, `Negative-1`, and `Negative-Random`. `Positive-MANE` sites (blue) are selected from the MANE database and supported by at least 100 alignments, while `Positive-Alt` sites (green) are present in the RefSeq database but missing from MANE, and also supported by at least 100 alignments. `Negative-1` sites (orange) occur on the opposite strand of a known gene and are supported by only 1 alignment, and Negative Random sites (red) are random GT-AG pairs on the opposite strand that do not overlap with any known splice sites and have no alignment support. **(b)** DNA logos for each type of splice site show the conservation of residues at each position in the training set created for this study. Each panel displays 40 nucleotides, centering on the donor and acceptor sites. The y-axis represents the level of conservation, measured in bits, with 0 representing equal probability among “A”, “C”, “G”, and “T”, and 2 representing a single nucleotide dominating the position.

#### 4.1.1 Positive dataset creation

We first ran TieBrush ^35^ to consolidate the HISAT2 alignments of 9,795 samples, collected across 31 histological types by the GTEx consortium project, into a single compressed BAM file. This TieBrush file was recently used to build the CHESS (v3) gene catalog ^32^. Following the consolidation, we extracted all splice junctions from the BAM file using the Regtools `junctions extract` command ^36^.

To increase the accuracy of the donors and acceptors extracted from the TieBrush file, we filtered the junctions to retain only those that were supported by more than 100 alignments. We used the RefSeq annotation to extract the coordinates of all protein-coding genes and intersected these with the reliable (100+ alignments) splice sites. The resulting splice sites were categorized as “`Positive-MANE`” if they were in MANE, and “`Positive-Alt`” if they were only in RefSeq but not in MANE. This gave us a set of annotated splice junctions from protein-coding genes that also had strong alignment evidence. Overall, we identified 180,195 splice sites in `Positive-MANE` and 85,908 splice sites in `Positive-Alt`.

#### 4.1.2 Negative dataset creation

We considered several ways to generate negative splice junctions. For instance, we could select random unannotated GT-AG pairs from the genome as negative splice sites. GT-AG pairs account for more than 98% of all splice sites in mammalian genomes in GenBank ^37^, but the vast majority of random GT-AG pairs are not splice junctions. However, the patterns of these artificial junctions might differ greatly from true splice sites, making them easy to learn but less useful for rejecting transcriptional noise. Arbitrarily labeling unannotated junctions as negatives would also be inappropriate because some of them may actually be spliced. This could happen because of incomplete annotation, transcriptional noise, or other reasons ^13, 38^.

Nonetheless, we needed to generate an accurate set of negative examples for training Splam, and we adopted two novel approaches. The first approach involved selecting random GT-AG pairs on the genome and requiring them to be on the opposite strand from known protein-coding genes. Overlapping genes are very rare in eukaryotes ^39, 40, 41, 42, 43^, so these will almost certainly be negative examples. We selected random GT-AG pairs in this fashion to generate 4,467,910 splice junctions, which we call the `Negative-Random` set.

Second, we wanted a set of negative examples that resembled true splice junctions, but that were likely to be non-functional. To generate this set, we chose splice junctions with only a single alignment supporting them in the TieBrush file and further required that they be on the strand opposite from a protein-coding gene. Because they have an alignment supporting them, these splice junctions might represent transcriptional noise, making them more useful for training. This approach generated a set of 2,486,305 splice junctions, which we refer to as `Negative-1`.

#### 4.1.3 Evaluating the donor and acceptor sites in positive and negative datasets

Figure 6b shows DNA logos for each of our positive and negative datasets. The donor and acceptor patterns for `Positive-MANE` and `Positive-Alt` are almost identical, indicating that splice junctions in MANE and alternatively spliced transcripts from RefSeq have the same consensus, as expected for positive examples. The two negative datasets show almost no pattern of conservation, although the `Negative-1` group has a very small amount of conservation immediately adjacent to each splice junction. The two groups of negatively labeled splice junctions are clearly distinguishable from the annotated true splice sites. <u>Table S12</u> reports the numbers of dinucleotides at the donor and acceptor sites, and <u>Figure S10</u> illustrates the ratio of canonical to non-canonical splice sites.

### 4.2 Splam input and output data encoding

For curating input splice sites, we extract a 400nt sequence centered around each end of an intron, learning the splicing pattern at the junction level. This approach facilitates local prediction of a splicing event, removes any canonical transcript bias, and eliminates the reliance on a larger window size, which for SpliceAI includes the whole intron and 10,000nt of flanking sequences.

For every splice junction coordinate, we used gffread ^44^ to extract sequences of 400nt around donors and acceptor sites, in total 800nt. In cases where the intron length falls between 200 and 400 base pairs, we select a 200nt sequence from both the donor and acceptor sides, with overlapping sequences allowed. For introns shorter than 200nt, we include the entire intron with Ns inserted to extend the sequence to 200nt for both donor and acceptor sites. The input sequences are then one-hot encoded, with [*A*, *C*, *G*, *T*, *N*] represented as [1, 0, 0, 0], [0, 1, 0, 0], [0, 0, 1, 0], [0, 0, 0, 1], and [0, 0, 0, 0].

In terms of the labeling of splice sites, we one-hot encoded the 200^th^ nucleotide (the donor site) as [0, 0, 1], and the 600^th^ nucleotide (the acceptor site) as [0, 1, 0] (using a 0-based numbering system). Nucleotides at non-donor and acceptor sites were labeled as [0, 0, 0]. For the input sequences, over 95% of them had GT as their 200^th^ and 201^st^ nucleotides, and AG as their 598^th^ and 599^th^ nucleotides. We also note that 8 donor sites had ‘TT’ and another 8 donor sites had ‘GA’ in the MANE database (<u>Table</u> <u>S12</u>).

### 4.3 Splam model architecture design

The model design of Splam mainly follows the architecture of SpliceAI with some optimizations. We utilized a deep residual convolutional neural network (CNN) that incorporates grouped dilated convolution layers within the residual units (Figure 1a). The input to the model is 800nt of one-hot encoded DNA sequence, and the output is the probability for each base if it is a donor site, acceptor site, or neither in the dimension of 3 × 800 (<u>Methods 4.2</u>). The three scores are constrained to sum to 1. We constructed the Splam model using PyTorch framework version 1.13.0 ^45^.

The Splam model consists of 20 residual units, each containing two convolutional layers, and each convolutional layer follows a batch normalization and a Leaky rectified linear unit (LReLU) ^46^ with a negative slope of 0.1 as shown in Figure 1a. We define the inputs of a residual unit using four hyperparameters: *F* for the number of filters, *W* for the window size, *D* for the dilation rate, and *G* for the number of groups. These input hyperparameters are represented as (*F*, *W*, *D*, *G*) in Figure 1b. The input to a convolutional layer is defined as (*F*, *W*, *D*). Two key ideas used in the Splam model architecture are (1) residual connection and (2) grouped dilated convolution.

Residual units were first proposed by He et al. in 2015 ^47^ to solve the training accuracy degradation problem ^48, 49^. The residual mappings (shortcut connections) allow one to train the deeper model by simple stochastic gradient descent (SGD) with backpropagation and can result in accuracy gains from the increment in the depth of the model. The concept of residual connection is widely used in deep neural networks today to prevent the vanishing/exploding gradient problem. In Splam, the input of a residual unit is connected to its output through a residual connection.

In the Splam model, the 1-D convolutional layers in each residual unit have a dilation rate parameter, denoted as *D*. The dilation rate determines the spacing between the values in the convolutional filter, allowing the network to have a larger receptive field without increasing the number of parameters without loss of resolution or coverage ^50^. The dilated convolution operation is defined as follows:

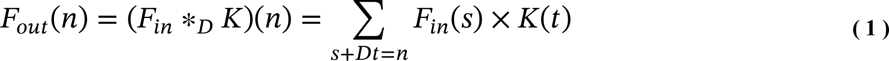

where *F*_*in*_ is the input feature, *F_out_* is the out feature, *K* is the convolutional kernel, *n* represents the index of the output feature, and ∗_*D*_ represents the dilated convolution operation (Equation ( 1 )). In Splam, the dilated convolutional kernel captures features from a neighboring region spanning *D* × (*W* − 1) in *F*_*in*_, encompassing a total of 2*D* × (*W* − 1) neighboring positions for the input of the respective residual unit.

The idea of grouped convolution in Splam was inspired by the cardinality concept, which refers to the number of parallel paths present in a block, as seen in ResNext ^51^. The concept of grouped convolution dates back to AlextNet ^19^, and is implemented in widely used open-source machine learning frameworks such as Caffe ^52^, Keras ^53^, TensorFlow ^54^, and PyTorch ^45^. The use of grouped convolution allows for substantial memory savings with minimal loss of accuracy, which can further push the model deeper ^55^. In a grouped convolution with *G* groups, *F*/*G* filters are applied to each *F*/*G* portion of the input, resulting in a *G* × reduction in the number of parameters used. In Splam, each residual unit consists of two convolutional layers where *F* is set to 64 and the group parameter is set to 4.

A group of four residual units (RUs) forms a bigger residual group, and 20 RUs are clustered into five residual groups, in the order of blue, green, orange, yellow, and purple in Figure 1. Residual groups are stacked such that the output of the *i*^th^ residual group is forward-connected to the *i* + 1^th^ residual group (forward connections colored in blue).

Furthermore, the output of each residual group undergoes a convolutional layer, with the parameters (64, 1, 1), which is then added to all the other outputs of residual groups (residual connections colored in red), which then is passed into the last convolutional layer in (3, 1, 1) and a softmax layer, generating a 3- dimensional input Tensor with three channels lying in the range [0, 1] and summing to 1. *F* is set to 64 for all convolutional layers. For each residual group, *W* is set to 11, 11, 11, 21, and 21, and *D* is set to 1, 5, 10, 15, and 20, in sequence. *G* is 1 by default for all convolutional layers but set to 4 in the residual units. For each nucleotide position, its total neighboring span in the Splam model is 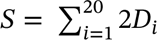 × (*W*_*i*_ − 1). In total, there are 651,715 parameters in Splam.

### 4.4 Model training and testing

Splam was trained on a MacBook Pro 2021 Apple M1 Pro chip with 16 GPU cores, 16 Neural Engine cores, and 16 GB unified RAM using PyTorch “mps” mode.

#### 4.4.1 Splitting splice junctions into training and testing datasets

After collecting and curating the splice junctions to be used for training and testing, we divided them into two datasets: one for model training and the other for testing. For model training, we utilized all the splice sites on the main chromosomes, except Chromosome 1 and 9. For model testing, we used the splice sites on the held-out Chromosomes 1 and 9, after removing splice sites from genes that had paralogs on the other chromosomes.

As discussed in the main text, we curated two sets of positive splice sites: `Positive-MANE` and `Positive-Alt`. Out of the 180,195 splice sites in `Positive-MANE`, 154,249 were allocated to the training dataset and 25,946 to the testing dataset. For `Positive-Alt`, out of the 85,908 splice sites, 73,789 were allocated to the training dataset and 12,119 to the testing dataset. In total, 228,038 positive splice sites were used for model training. We also curated two types of negative splice sites, Negative Random and `Negative-1`. Out of the 4,467,910 splice sites in `Negative-Random`, 3,834,401 were allocated to the training dataset and 633,509 to the testing dataset. For `Negative-1`, out of the 2,486,305 splice sites, 2,134,002 were allocated to the training dataset and 352,303 to the testing dataset. In total, we randomly selected three times as many negative splice sites as positive splice sites, resulting in 684,114 negative splice sites being used for model training. The positive to negative weighting in the loss function is three to one.

#### 4.4.2 Four test datasets for evaluating Splam

We generated four test datasets, namely Test40K, Test22K-MANE, Test22K-Alt, and Test24K. These datasets were derived from sampling the human splice junctions test dataset described in <u>Methods</u>.

Test40K encompasses 10,000 splice junctions each from `Positive-MANE`, `Positive-Alt`, `Negative-1`, and `Negative-Random` examples, maintaining a positive-to-negative ratio of one-to-one.

Test22K-MANE, Test22K-Alt, and Test24K share the same 10,000 `Negative-1` and 10,000 `Negative-Random` examples with Test40K. To simulate the sparsity of true junctions in the genome, we created a test set with a positive-to-negative ratio of 1:10, downsampling the initial 10,000 `Positive-MANE` and 10,000 `Positive-Alt to` 2,000 `Positive-MANE` and 2,000 `Positive-Alt,` respectively.

In sum, Test22K-MANE consists of 2,000 `Positive-MANE,` 10,000 `Negative-1`, and 10,000 `Negative-Random` examples; Test22K-Alt consists of 2,000 `Positive-Alt,` 10,000 `Negative-1`, and 10,000 `Negative-Random` examples; and Test24K consists of 2,000 `Positive-MANE`, 2,000 `Positive-Alt,` 10,000 `Negative-1`, and 10,000 `Negative-Random`. We used these datasets to evaluate Splam’s ability to recognize splice junctions discussed in <u>Results 2.1</u>.

#### 4.4.3 Splam training hyperparameters

To train Splam, we used a batch size of 100 and trained it for 15 epochs. We employed the AdamW ^56^ optimizer with a default learning rate of 0.03. A 1000-step warmup was utilized ^57^, with the learning rate increasing linearly from 0 to 0.03. The learning rate then decreased following the values of the cosine function between 0.03 to 0 ^58^.

We further improved Splam’s performance by changing the loss function. Instead of the commonly-used cross entropy (Equation ( 2 )), we replaced it with focal loss (Equation ( 3 )) ^59^.

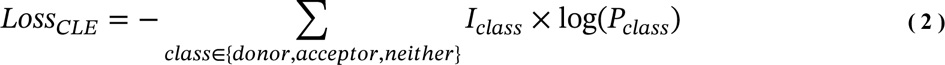

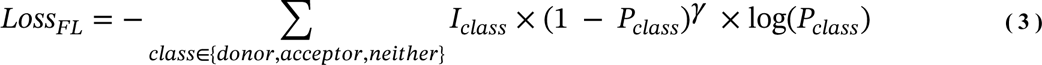

*I_class_* is the ground-truth label and *P*_*class*_ is the Splam output score for each class (donor, acceptor, or neither). Comparing the two equations, we can see that Equation ( 3 ) has an additional modulating term, (1 − *P*_*class*_)^*γ*^. This term scales the cross-entropy loss dynamically, with the scaling factor decreasing to zero as confidence in the correct class increases. In other words, the focal loss approach puts more emphasis on the misclassified challenging data points where Splam is more likely to make incorrect predictions and penalizes these data points by an additional (1 − *P*_*class*_)^*γ*^ scale. In Splam, we set *γ* = 2. In summary, focal loss quantifies the level of inaccuracy in predictions by down-weighting the influence of easy examples during training, enabling the model to swiftly prioritize challenging examples.

#### 4.4.4 Paralogs removal in testing dataset

We removed the paralogs from the testing dataset as follows: we first split the transcripts on Chromosome 1 and 9 and the remaining chromosomes and extracted the transcript sequences using gffread ^44^ into two FASTA files. We then used nucmer ^60^ with default parameters to align the two files and ran `show-coords` with the `-lcr` option to obtain the alignment coordinates. We removed alignments with sequence similarity greater than 80% and the alignment coverage percentage for the query greater than 50%. We found a total of 2,943 paralogous transcripts, and any splice sites in the testing dataset that were from these paralogs were removed.

For Splam model testing, we randomly selected 10,000 splice sites from the paralog-removed testing dataset for each of the four test set groups.

#### 4.4.5 Testing SpliceAI

In order to achieve optimal results with SpliceAI, the final splice site score was determined by averaging the outputs of five models, as recommended by the developers of SpliceAI. Throughout our experiments, we employ this ensemble approach for all SpliceAI analyses.

### 4.5 Running Splam on closely-related and distant non-human species

In this experiment, we aimed to evaluate Splam’s ability to generalize to different species by benchmarking the performance of Splam against SpliceAI in scoring donor-acceptor site pairs. Thus, we took the same approach as described in <u>Results 2.1</u> and ran both programs at the splice junction level: collecting junctions from non-human species and evaluating one junction at a time.

#### 4.5.1 Extracting splice junctions from GFF files as the positive testing dataset

We downloaded the target non-human reference annotations in GFF format, along with assembly reports and genome sequences in FASTA format, from the NCBI FTP server (specifically, *Pan troglodytes* and *Mus musculus*) and TAIR database (*Arabidopsis thaliana*) at https://www.arabidopsis.org/. For each genome, we filtered out any incorrectly overlapping exons and extracted introns from all isoforms of each gene as splice junctions using gffutils version 0.12.0 ^53^. The final output of this step is a BED file with supporting strand, parent ID, and chromosome information. This comprises the positive dataset to be used for evaluation.

#### 4.5.2 Creating the negative testing dataset

To produce compelling pseudo-splice-junction data for the negative dataset, we compiled the protein-coding genes for each species and followed a similar approach as in the creation of the human `Negative-Random` dataset (<u>Methods 4.1.2</u>). For each gene, we selected the strand opposite to the given protein-coding gene locus. Second, we generated a random start position within the boundaries of the locus and searched forward until finding a canonical GT donor site if on the + strand, or CT acceptor site if on the − strand. If none were found, we ended the current attempt. Next, we selected a random starting intron length from the interval [200, 20000], and jumped forward by this amount from the initial site. Starting from here, we searched forward to find a canonical AG acceptor site if on the + strand, or AC donor site if on the − strand. If none were found, we ended the current attempt. Otherwise, we recorded the donor site and acceptor site positions as a splice junction. We repeated this process for a total of 20 attempts per gene locus to produce our negative dataset.

#### 4.5.3 Processing splice junctions for Splam and SpliceAI

Using the obtained coordinates from the positive and negative datasets, we pre-processed the splice junctions to fit the requirements for each model. For Splam, we extracted a 400nt uppercase sequence for both donor and acceptor sites, as described in <u>Method 4.2</u>, resulting in an 800nt input sequence for each splice junction. In cases where the donor and acceptor sites were <400nt apart, we allowed the flanking sequences to overlap in the intronic region, resulting in duplicated segments near the center of the input sequence. If the sites were exactly 200nt apart, the intronic flanking sequences of the two sites would be identical. In cases where the sites were <200nt apart, we truncated the flanking sequences to the end of the intronic region (i.e. the donor or acceptor site itself) and padded the remainder with Ns. No matter the case, the resulting input sequence would always be exactly 800nt long.

For SpliceAI, we extracted the entire intron sequence with a 10,400nt context (5,200nt flanking each side) as described in <u>Method 4.8</u>. If the flanking sequence went out of the chromosome’s range, we truncated to the boundary of the chromosome and padded the remainder with Ns. Thus, the length of the SpliceAI input sequence was 10.4Kb plus the intron length.

In order to run the SpliceAI models in a reasonable time, we randomly sampled 25,000 splice junctions from both the positive and negative datasets, and ran Splam and SpliceAI on each junction, following the protocols outlined in Method 4.8.

### 4.6 Applying Splam to remove spurious spliced alignments

One application of Splam is to remove spliced alignments with low-scoring splice junctions. Users can run Splam on alignment files (BAM). We assess the results at the intron level and the transcriptome assembly level.

#### 4.6.1 Sample information of RNA-Seq data

We randomly selected 20 RNA-seq samples that were collected and sequenced by the Lieber Institute for Brain Development. Among these samples, 10 were prepared using poly-A capture and 10 were prepared using ribosomal RNA depletion. These samples were obtained by the Lieber Institute from developing and mature human dorsolateral prefrontal cortex (DLPFC). All FASTQ files of these samples are publicly available via Globus collections jhpce#bsp2-dlpfc at https://app.globus.org/file-manager?origin_id=0dd03924-6853-11e9-bf44-0e4a062367b8&origin_path=%2F and <u>jhpce#bsp2-hippo</u> at https://app.globus.org/file-manager?origin_id=96be20a2-6853-11e9-bf44-0e4a062367b8&origin_path=%2F. We then aligned these samples to GRCh38 version 40 patch 14 with HISAT2.

#### 4.6.2 Splam removal procedure workflow overview

Splam removal procedure would occur as a step in RNA-Seq data analysis after spliced alignment and before transcriptome assembly (Figure 5a). The Splam removal step is highlighted by the red box, and a detailed breakdown of the trimming process is presented in Figure 5b.

Before running the Splam spliced alignment removal procedure, users need to prepare three files, which are (1) a target alignment file in BAM format, (2) a reference genome in FASTA format, and (3) the released Splam model in PT format. This workflow can be done in three lines of code:

1. Extracting splice junctions from the alignment file:

~~~
$ splam extract −P −o tmp_out <INPUT alignment file>
~~~
2. Scoring extracted splice junctions:

~~~
$ splam score −G <INPUT reference genome> −m <INPUT splam model> −o tmp_out tmp_out/junction.bed
~~~

In this step, Splam takes the splice junctions in the BED file generated in the previous step, encodes the data described in <u>Methods 4.2</u>, and scores them using the trained Splam model (see <u>Methods 4.3</u> and <u>4.4</u>). Splam automatically detects the environment and runs in “`cudo`” mode if CUDA is available. If the computer is running macOS, Splam will check if `mps` mode is available. If neither “`cuda`” nor “`mps`” are available, Splam will run in “`cpu`” mode.

3. Cleaning up the alignment file:

~~~
$ splam clean −o tmp_out
~~~

Subsequently, Splam proceeds to iterate through the alignments. Splam can run on alignment files of both single-end and paired-end RNA-Seq samples. Any alignment containing any spurious splice junctions is removed, and if it is paired, Splam updates the flags to unpair reads for both the aligned read and its mate. Moreover, if a read is multi-mapped, Splam updates the NH tag for the alignment itself and all other alignments. The outcome of this spliced alignment removal step is a cleaned, sorted alignment file (BAM) along with an alignment file comprising the discarded alignments.

#### 4.6.3 Metrics for intron matching

Two metrics, intron precision and recall, are used to assess the alignment file at the intron level. Intron precision is calculated as the ratio of true positives (TP) to the sum of true positives and false positives (TP + FP), where TP represents the number of introns in the alignment file that match the RefSeq annotation, while FP represents the number of introns in the alignment file that do not match the annotation. Intron recall, on the other hand, is computed as TP/(TP + FN). In this case, TP again represents the number of introns in both the alignment and the RefSeq annotation, while FN denotes the number of introns present in the annotation file but not in the alignment file.

#### 4.6.4 Metrics for transcript assembly

In evaluating the assembled transcripts, we examined two metrics: the total number of assembled transcripts that match the RefSeq database and the transcript precision. Transcript precision is calculated as the ratio of true positives (TP) to the sum of true positives and false positives (TP + FP). Here, TP represents the assembled matching transcripts, while FP represents the hypothetical transcripts. The transcript precision reflects the proportion of assembled transcripts that are actually inside the annotation file.

### 4.7 Applying Splam to score introns in annotation files and assembled transcripts

Users can run Splam on (1) annotation files or (2) assembled transcripts in three lines of code. Splam iterates through the GFF file, extracts all introns in transcripts, and writes their coordinates into a BED file. The BED file consists of six columns: CHROM, START, END, JUNC NAME, INTRON NUM, and STRAND. Splam scores every extracted intron and outputs the scores of each donor and acceptor site.

$ splam extract −o tmp_out <INPUT annotation file>

$ splam score −G <INPUT reference genome> −m <INPUT Splam model> −o tmp_out tmp_out/junction.bed

Last, users can generate reports on the number of low-scoring splice junctions in each transcript and filter out transcripts with a high ratio of bad splice junctions using a user-defined threshold.

$ splam clean −o tmp_out

### 4.8 SpliceAI benchmarking

All the experiments of SpliceAI are conducted on a 24-core, 48-thread Intel(R) Xeon(R) Gold 6248R Linux computer with 1024 GB memory, running in “cpu” mode and using a single thread of execution.

#### 4.8.1 Formatting input data

The SpliceAI model requires a 10,000nt context to run, meaning that the input intron sequence needs to be padded with a 5,000nt flanking sequence on either side. We added an additional 200nt to both flanking sequences to better compare with Splam and prevent donor and avoid the donor and acceptor sites from being the boundary nucleotides. In the “SpliceAI” version, we extracted the full 5.2Kb flanking sequence from the genome. In the “SpliceAI-10k-Ns” version, we replaced the outer 5,000nt on each flanking sequence with Ns (zero-padding, as outlined in ^23^), leaving only a 200nt sequence from the genome.

#### 4.8.2 Testing the SpliceAI model

Pursuant to the methods described in ^23^, we ran SpliceAI on large datasets in discrete batches, with a batch size of 500 splice junctions. We one-hot encoded the input sequence, loaded the resulting Tensor into the SpliceAI model, and extracted the output into donor, acceptor, and non-splice-site channels. Each channel was a list containing the corresponding scores for every nucleotide in the input sequence. The channels were then written to their respective files, with every line representing a different splice junction. We repeated this process for all 5 SpliceAI models, then averaged the donor and acceptor site scores.

#### 4.8.3 Executing and speeding up SpliceAI predictions

In order to do batch prediction, we created a separate bash script to execute the above model testing process in increments of 500 (batch size), up to the size of the input dataset. We employed a wrapper script for ease of access and to prevent memory leakage.

Additionally, to speed up the pipeline across models and for both “noN” and “N” modes, we multi-threaded the bash execution to run the 10 targets (5 models, 2 modes) concurrently.

### 5. Data and code availability

The Splam project is freely available on github at: <u>github.com/Kuanhao-Chao/splam</u>, and is available on PyPi: https://pypi.org/project/splam/. The Splam documentation is available at: <u>ccb.jhu.edu/splam</u>.

Splam was built and trained using the PyTorch ^45^ framework. The core of Splam is implemented in C++ using hstlib ^61^ and the samtools merge and sort scripts ^62^. C++ libraries were compiled and linked to Python using Pybind11 ^63^ (https://github.com/pybind/pybind11). The scripts for Splam training and data analysis are freely available on GitHub at: <u>github.com/Kuanhao-Chao/splam-analysis-results</u>.

## 6. Acknowledgments

We thank Markus J. Sommer, Jakob Heinz, and Natalia Rincon for their help in naming Splam.

## 7. Funding

This research was supported in part by the U.S. National Institutes of Health under grants R0I-HG006677 and R01-MH123567, and by the U.S. National Science Foundation under grant DBI-1759518.

## 8. Authors’ contributions

KC, SLS, and MP designed the research. KC designed and trained the Splam model. KC ran all experiments on human splice sites. AM and KC conducted the cross-species generalization analysis. KC developed the software to improve alignment files. KC, AM, SLS, and MP wrote the manuscript.

## Notes

http://ccb.jhu.edu/splam/

https://github.com/Kuanhao-Chao/splam

### Competing Interest Statement

The authors have declared no competing interest.

### Summary of Updates

We update some grammar errors in the manuscript.

## References

1. Berget, S.M., Moore, C. & Sharp, P.A. Spliced segments at the 5′ terminus of adenovirus 2 late mRNA. Proceedings of the National Academy of Sciences 74, 3171–3175 (1977).

2. Baralle, F.E. & Giudice, J. Alternative splicing as a regulator of development and tissue identity. Nature reviews Molecular cell biology 18, 437–451 (2017).

3. Johnson, J.M. et al. Genome-wide survey of human alternative pre-mRNA splicing with exon junction microarrays. Science 302, 2141–2144 (2003).

4. Kalsotra, A. & Cooper, T.A. Functional consequences of developmentally regulated alternative splicing. Nature Reviews Genetics 12, 715–729 (2011).

5. Licatalosi, D.D. & Darnell, R.B. RNA processing and its regulation: global insights into biological networks. Nature Reviews Genetics 11, 75–87 (2010).

6. Kornblihtt, A.R. et al. Alternative splicing: a pivotal step between eukaryotic transcription and translation. Nature reviews Molecular cell biology 14, 153–165 (2013).

7. Pan, Q., Shai, O., Lee, L.J., Frey, B.J. & Blencowe, B.J. Deep surveying of alternative splicing complexity in the human transcriptome by high-throughput sequencing. Nature genetics 40, 1413–1415 (2008).

8. Trapnell, C., Pachter, L. & Salzberg, S.L. TopHat: discovering splice junctions with RNA-Seq. Bioinformatics 25, 1105–1111 (2009).

9. Au, K.F., Jiang, H., Lin, L., Xing, Y. & Wong, W.H. Detection of splice junctions from paired-end RNA-seq data by SpliceMap. Nucleic acids research 38, 4570–4578 (2010).

10. Wang, K. et al. MapSplice: accurate mapping of RNA-seq reads for splice junction discovery. Nucleic acids research 38, e178–e178 (2010).

11. Dobin, A. et al. STAR: ultrafast universal RNA-seq aligner. Bioinformatics 29, 15–21 (2013).

12. Kim, D., Paggi, J.M., Park, C., Bennett, C. & Salzberg, S.L. Graph-based genome alignment and genotyping with HISAT2 and HISAT-genotype. Nature biotechnology 37, 907–915 (2019).

13. Varabyou, A., Salzberg, S.L. & Pertea, M. Effects of transcriptional noise on estimates of gene and transcript expression in RNA sequencing experiments. Genome research 31, 301–308 (2021).

14. Pertea, M., Lin, X. & Salzberg, S.L. GeneSplicer: a new computational method for splice site prediction. Nucleic acids research 29, 1185–1190 (2001).

15. Yeo, G. & Burge, C.B. in Proceedings of the seventh annual international conference on Research in computational molecular biology 322–331 (2003).

16. Dogan, R.I., Getoor, L., Wilbur, W.J. & Mount, S.M. SplicePort—an interactive splice-site analysis tool. Nucleic acids research 35, W285–W291 (2007).

17. Degroeve, S., Saeys, Y., De Baets, B., Rouzé, P. & Van de Peer, Y. SpliceMachine: predicting splice sites from high-dimensional local context representations. Bioinformatics 21, 1332–1338 (2005).

18. Sonnenburg, S., Schweikert, G., Philips, P., Behr, J. & Rätsch, G. in BMC bioinformatics, Vol. 8 1–16 (BioMed Central, 2007).

19. Krizhevsky, A., Sutskever, I. & Hinton, G.E. Imagenet classification with deep convolutional neural networks. Advances in neural information processing systems 25 (2012).

20. Dai, J., Li, Y., He, K. & Sun, J. R-fcn: Object detection via region-based fully convolutional networks. Advances in neural information processing systems 29 (2016).

21. Chen, L.-C., Papandreou, G., Kokkinos, I., Murphy, K. & Yuille, A.L. Deeplab: Semantic image segmentation with deep convolutional nets, atrous convolution, and fully connected crfs. IEEE transactions on pattern analysis and machine intelligence 40, 834–848 (2017).

22. Zeiler, M.D. & Fergus, R. in Computer Vision–ECCV 2014: 13th European Conference, Zurich, Switzerland, September 6–12, 2014, Proceedings, Part I 13 818-833 (Springer, 2014).

23. Jaganathan, K. et al. Predicting splicing from primary sequence with deep learning. Cell 176, 535–548. e524 (2019).

24. Wang, R., Wang, Z., Wang, J. & Li, S. SpliceFinder: ab initio prediction of splice sites using convolutional neural network. BMC bioinformatics 20, 1–13 (2019).

25. Scalzitti, N. et al. Spliceator: Multi-species splice site prediction using convolutional neural networks. BMC bioinformatics 22, 1–26 (2021).

26. Zuallaert, J. et al. SpliceRover: interpretable convolutional neural networks for improved splice site prediction. Bioinformatics 34, 4180–4188 (2018).

27. Vaswani, A. et al. Attention is all you need. Advances in neural information processing systems 30 (2017).

28. Ji, Y., Zhou, Z., Liu, H. & Davuluri, R.V. DNABERT: pre-trained Bidirectional Encoder Representations from Transformers model for DNA-language in genome. Bioinformatics 37, 2112–2120 (2021).

29. Dalla-Torre, H. et al. The Nucleotide Transformer: Building and Evaluating Robust Foundation Models for Human Genomics. bioRxiv, 2023.2001. 2011.523679 (2023).

30. Morales, J. et al. A joint NCBI and EMBL-EBI transcript set for clinical genomics and research. Nature 604, 310–315 (2022).

31. O’Leary, N.A. et al. Reference sequence (RefSeq) database at NCBI: current status, taxonomic expansion, and functional annotation. Nucleic acids research 44, D733–D745 (2016).

32. Varabyou, A. et al. CHESS 3: an improved, comprehensive catalog of human genes and transcripts based on large-scale expression data, phylogenetic analysis, and protein structure. bioRxiv, 2022.2012. 2021.521274 (2022).

33. Pertea, M. et al. StringTie enables improved reconstruction of a transcriptome from RNA-seq reads. Nature biotechnology 33, 290–295 (2015).

34. Moore, M.J. Intron recognition comes of AGe. Nature structural biology 7, 14–16 (2000).

35. Varabyou, A., Pertea, G., Pockrandt, C. & Pertea, M. TieBrush: an efficient method for aggregating and summarizing mapped reads across large datasets. Bioinformatics 37, 3650-3651 (2021).

36. Feng, Y.-Y. et al. RegTools: Integrated analysis of genomic and transcriptomic data for discovery of splicing variants in cancer. BioRxiv 10, 436634 (2018).

37. Burset, M., Seledtsov, I.A. & Solovyev, V.V. Analysis of canonical and non-canonical splice sites in mammalian genomes. Nucleic acids research 28, 4364–4375 (2000).

38. Amaral, P., et al. The status of the human gene catalogue. arXiv preprint arXiv:2303.13996 (2023).

39. Loughran, G. et al. Unusually efficient CUG initiation of an overlapping reading frame in POLG mRNA yields novel protein POLGARF. Proceedings of the National Academy of Sciences 117, 24936–24946 (2020).

40. Pavesi, A. et al. Overlapping genes and the proteins they encode differ significantly in their sequence composition from non-overlapping genes. PloS one 13, e0202513 (2018).

41. Sanna, C.R., Li, W.-H. & Zhang, L. Overlapping genes in the human and mouse genomes. BMC Genomics 9, 169 (2008).

42. Veeramachaneni, V., Makalowski, W., Galdzicki, M., Sood, R. & Makalowska, I. Mammalian overlapping genes: the comparative perspective. Genome research 14, 280–286 (2004).

43. Wright, B.W., Molloy, M.P. & Jaschke, P.R. Overlapping genes in natural and engineered genomes. Nature Reviews Genetics 23, 154–168 (2022).

44. Pertea, G. & Pertea, M. GFF utilities: GffRead and GffCompare. F1000Research 9 (2020).

45. Paszke, A., et al. Automatic differentiation in pytorch. (2017).

46. Clevert, D.-A., Unterthiner, T. & Hochreiter, S. Fast and accurate deep network learning by exponential linear units (elus). arXiv preprint arXiv:1511.07289 (2015).

47. He, K., Zhang, X., Ren, S. & Sun, J. in Proceedings of the IEEE conference on computer vision and pattern recognition 770–778 (2016).

48. He, K. & Sun, J. in Proceedings of the IEEE conference on computer vision and pattern recognition 5353-5360 (2015).

49. Srivastava, R.K., Greff, K. & Schmidhuber, J. Highway networks. arXiv preprint arXiv:1505.00387 (2015).

50. Yu, F. & Koltun, V. Multi-scale context aggregation by dilated convolutions. arXiv preprint arXiv:1511.07122 (2015).

51. Xie, S., Girshick, R., Dollár, P., Tu, Z. & He, K. in Proceedings of the IEEE conference on computer vision and pattern recognition 1492-1500 (2017).

52. Jia, Y. et al. in Proceedings of the 22nd ACM international conference on Multimedia 675–678 (2014).

53. Dale, R. in GitHub repository, Vol. 2023 (https://github.com/daler/gffutils; 2017).

54. Abadi, M., et al. Tensorflow: Large-scale machine learning on heterogeneous distributed systems. arXiv preprint arXiv*:1603.04467* (2016).

55. Gibson, P., et al. in 2020 IEEE 31st International Conference on Application-specific Systems, Architectures and Processors (ASAP) 189–196 (IEEE, 2020).

56. Loshchilov, I. & Hutter, F. Fixing weight decay regularization in adam. (2018).

57. Goyal, P. et al. Accurate, large minibatch sgd: Training imagenet in 1 hour. arXiv preprint arXiv:1706.02677 (2017).

58. Loshchilov, I. & Hutter, F. Sgdr: Stochastic gradient descent with warm restarts. arXiv preprint arXiv*:1608.03983* (2016).

59. Lin, T.-Y., Goyal, P., Girshick, R., He, K. & Dollár, P. in Proceedings of the IEEE international conference on computer vision 2980-2988 (2017).

60. Marçais, G. et al. MUMmer4: A fast and versatile genome alignment system. PLoS computational biology 14, e1005944 (2018).

61. Bonfield, J.K. et al. HTSlib: C library for reading/writing high-throughput sequencing data. Gigascience 10, giab007 (2021).

62. Danecek, P. et al. Twelve years of SAMtools and BCFtools. Gigascience 10, giab008 (2021).

63. Moldovan, W.J.a.J.R.a.D. in GitHub repository, Vol. 2023 (https://github.com/pybind/pybind11; 2017).

