## Supplementary Figures and Tables for "Splam: a deep-learning-based splice site predictor that improves spliced alignments"

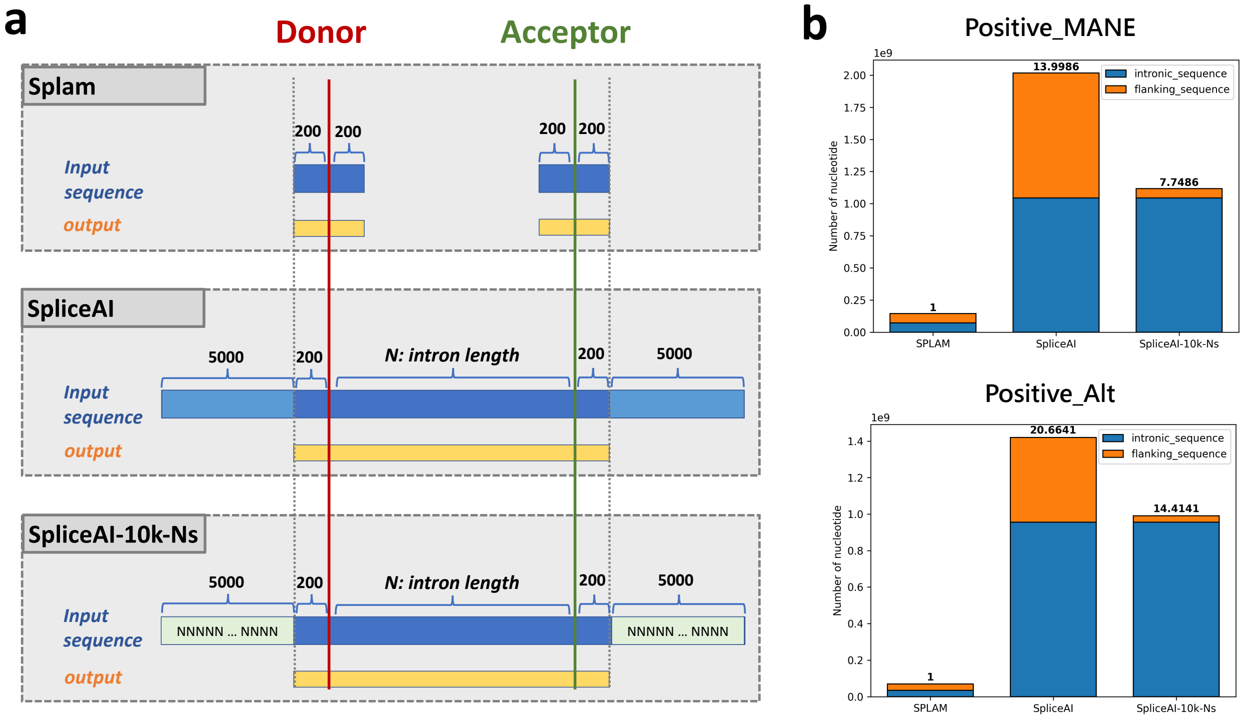


**Figure S1: (a)** The inputs and outputs for Splam, SpliceAI, and SpliceAI-10k-Ns for each splice junction. At the top is the input for Splam, with 400nt flanking the donor site and another 400nt flanking the acceptor site. The output is a set of labels for the 800nt, shown in yellow. The second row represents the input to SpliceAI in its standard configuration, with 200nt upstream and downstream of the donor and acceptor sites, the entire intron regardless of its length, and 10Kb of flanking sequence. The 200nt upstream and downstream sequences prevent the donor and acceptor sites from being the boundary nucleotides. The output of SpliceAI is similarly a set of labels for the region shown in yellow. The third row represents a slightly modified input designed for this study where the 5Kb flanking sequences are replaced with Ns, which we call SpliceAI-10k-Ns. **(b)** The amount of sequence used as input by Splam, SpliceAI, and SpliceAI-10k-Ns for the Positive-MANE and Positive-Alt datasets. Blue regions represent intronic sequences and orange regions represent flanking/exonic sequences. The ratio between the input size for Splam and SpliceAI is shown at the top of each bar. For Positive-MANE, the ratios of SpliceAI and SpliceAI-10k-Ns to Splam are 14.0 and 7.7, respectively. For Positive-Alt, the ratios of SpliceAI and SpliceAI-10k-Ns to Splam are 20.7, respectively.

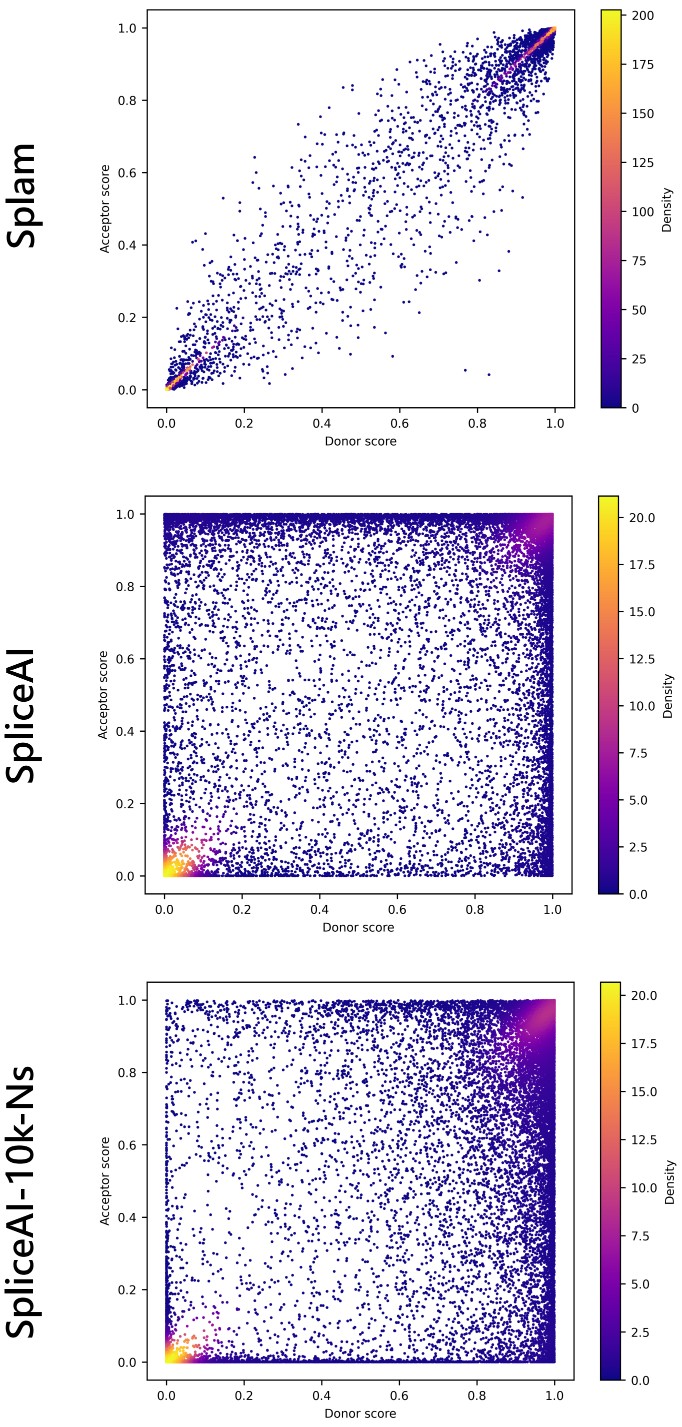


**Figure S2:** The splice junction score distribution for Splam, SpliceAI, and SpliceAI-10K-Ns. 10,000 splice junctions were randomly selected from the Positive-MANE, Positive-Alt, Negative-1, and Negative-Random test datasets, totaling 40,000 splice junctions. In the plot, each dot represents a splice junction, where the x-axis corresponds to the donor score and the y-axis represents the acceptor score. The dots are color-coded based on clustering density, with yellow indicating the highest density and blue indicating the lowest density.

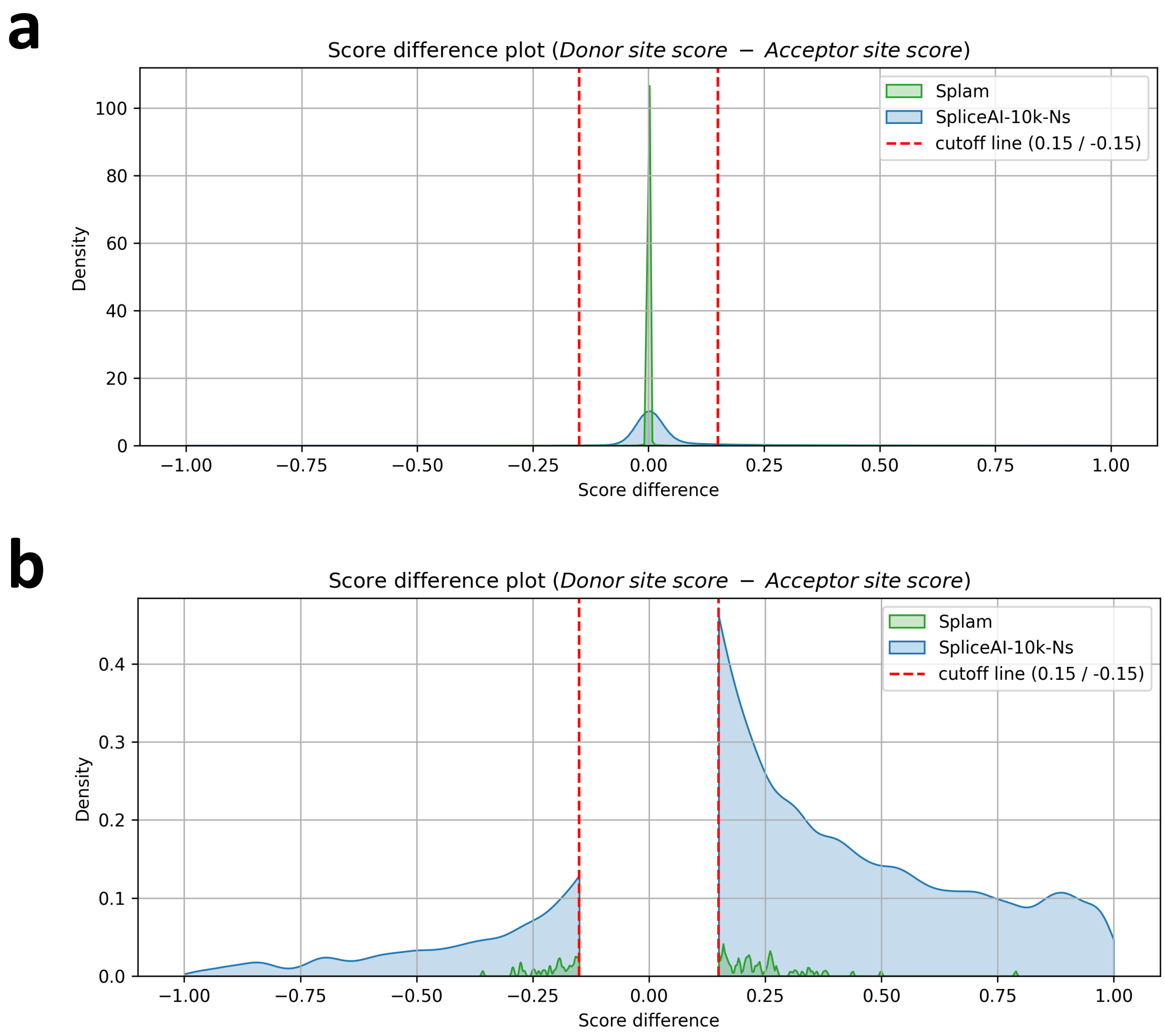


**Figure S3: (a)** Kernel density plots visualizing the differences between donor and acceptor scores (donor score − acceptor score). In **(b)** the score differences between -0.15 and 0.15 have been removed to provide a closer look at the splice junctions where the scores of donor and acceptor sites were relatively large. The green density plot represents Splam and the blue density plot represents SpliceAI-10k-Ns.

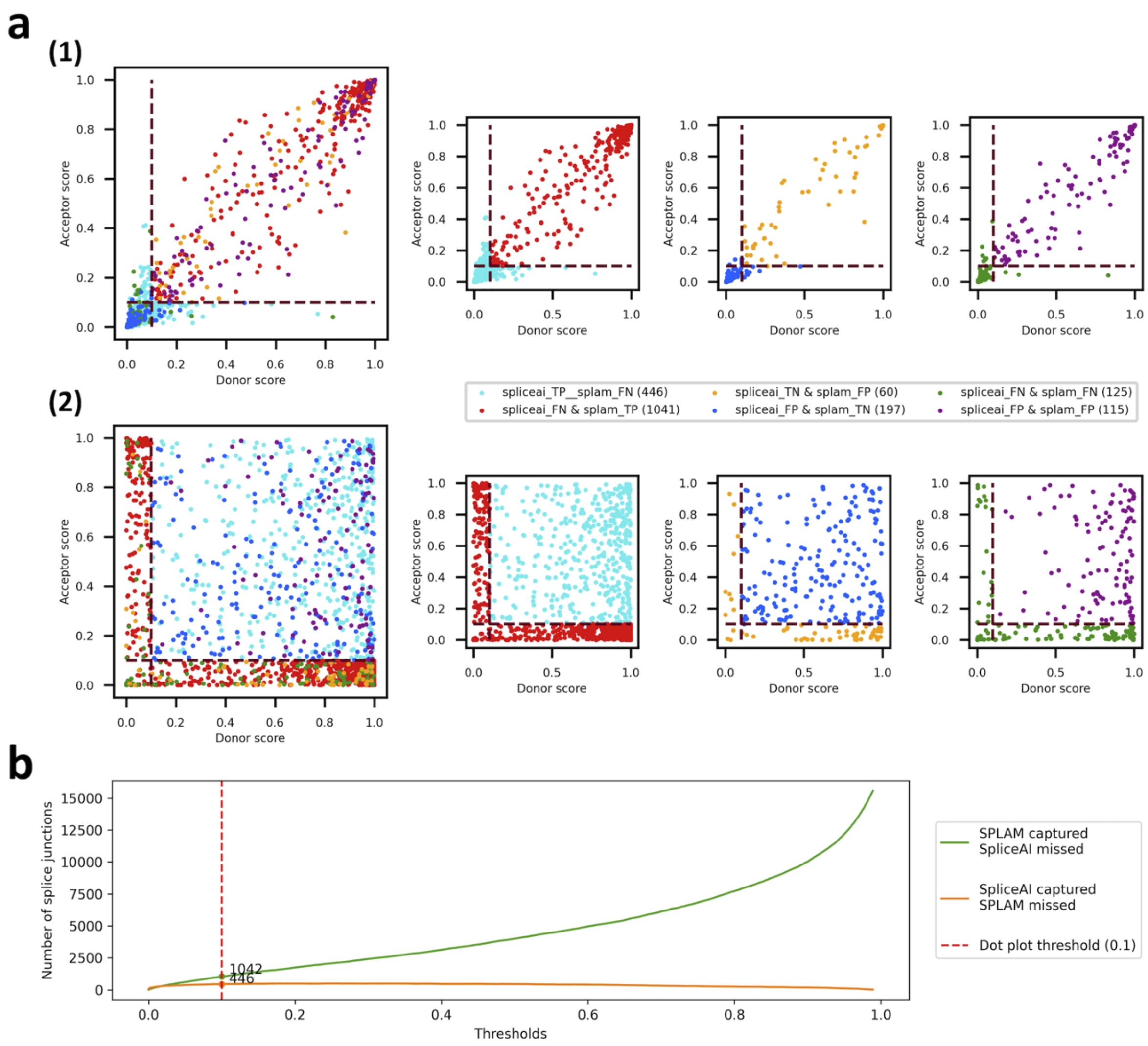


**Figure S4:** The results of comparing Splam and SpliceAI-10k-Ns. In Figure **(a)**, scatter plots **(1)** and **(2)** are depicted for Splam and SpliceAI-10k-Ns, where each dot represents a junction. The x-axis represents the donor score, while the y-axis represents the acceptor score. The red dashed lines indicate the cutoff threshold lines that determine whether a donor or acceptor is classified as good or bad. True positives (TPs), true negatives (TNs), false positives (FPs), and false negatives (FNs) are plotted based on the scores assigned by Splam **(1)** and SpliceAI **(2)**. The red dotted lines illustrate the thresholds used to label a point as true or false. To maintain clarity, these plots exclude the numerous points where both programs made correct predictions. The smaller subplots in the second and third columns display subsets of junctions where one program was correct while the other was incorrect (TP and FN, or FP and TN). The fourth column presents subsets of junctions where both programs made incorrect predictions (FP and FN). **(b)** The number of splice sites where Splam and SpliceAI disagree is shown. The green curve represents the instances where Splam correctly predicts them as true positives, but SpliceAI predicts them as false negatives. The orange curve represents the cases where SpliceAI correctly predicts them as true positives, but Splam predicts them as false negatives. The x-axis displays the threshold range from 0.001 to 0.999, while the y-axis represents the number of splice junctions. The vertical dashed red line signifies the 0.1 threshold, which is highlighted in **(a)**.

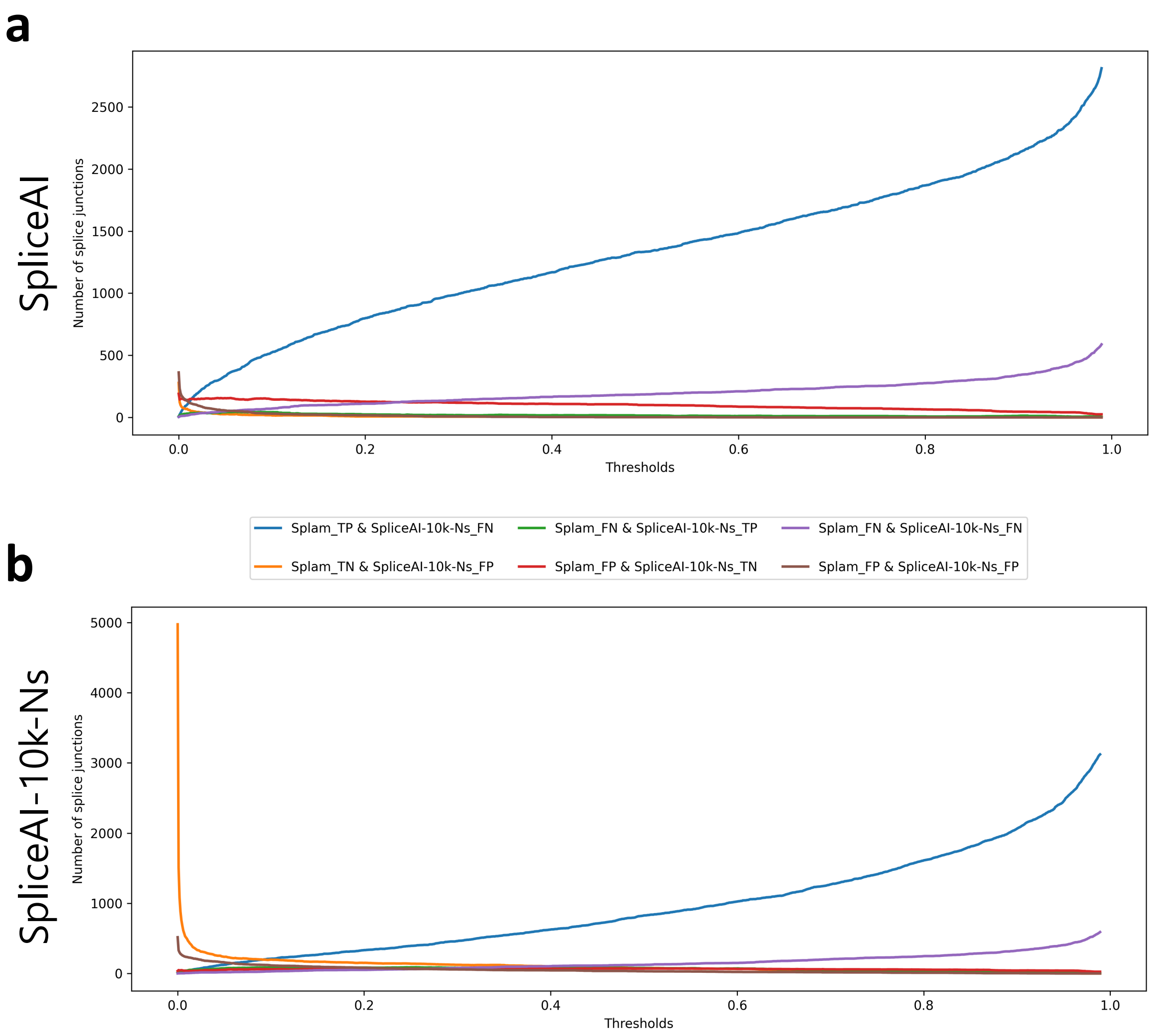


**Figure S5:** The number of splice junctions where Splam and SpliceAI disagree with each other or both programs get the wrong predictions. **(a)** The results comparing Splam and SpliceAI-10k. **(b)** The results comparing Splam and SpliceAI-10k-Ns. The blue curve represents the instances where Splam correctly predicts them as true positives, but SpliceAI predicts them as false negatives. The orange curve represents the cases where Splam correctly predicts them as true negatives, but SpliceAI predicts them as false positives. The green curve represents the cases where SpliceAI correctly predicts them as true positives, but Splam predicts them as false negatives. The red curve represents the cases where SpliceAI correctly predicts them as true negatives, but Splam predicts them as false positives. The purple curve represents both Splam and SpliceAI wrongly predicting them as false negatives, and the brown curve represents both Splam and SpliceAI wrongly predicting them as false positives. The x-axis displays the threshold range from 0.001 to 0.999, while the y-axis represents the number of splice junctions.

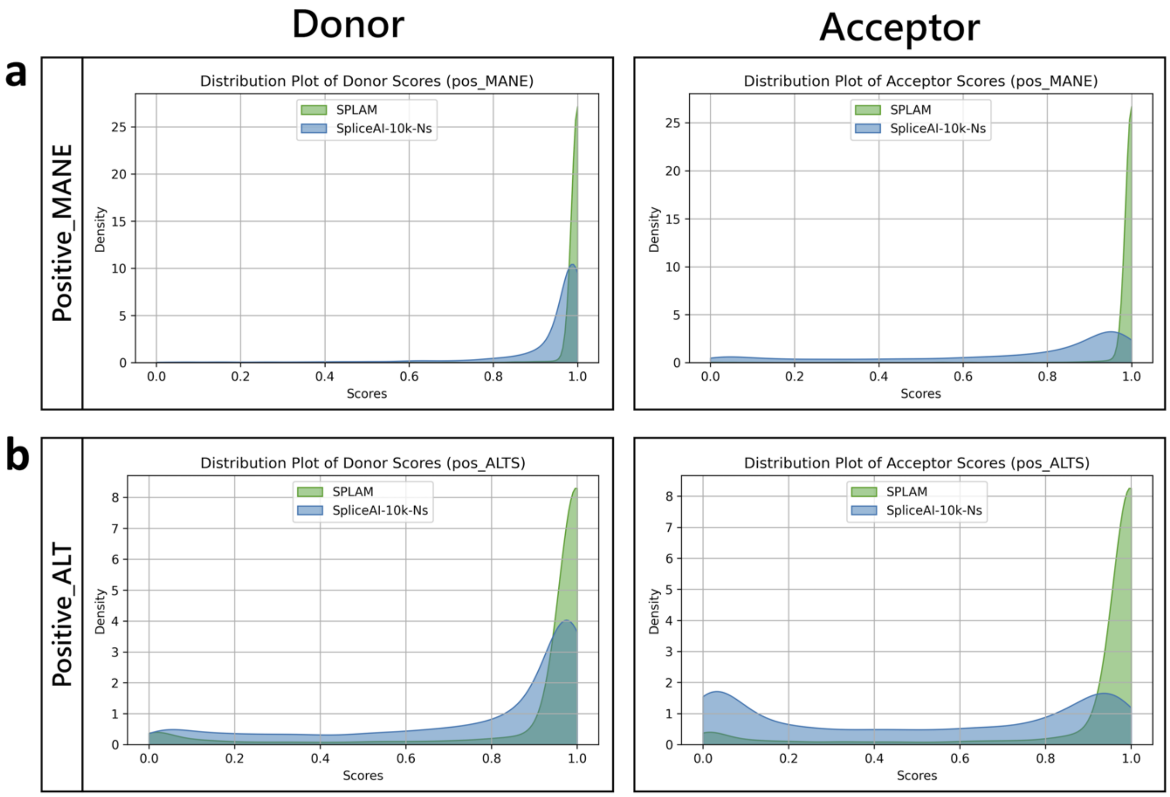


**Figure S6:** The score kernel density plots for Splam and SpliceAI-10k-Ns. The x-axis is the scores ranging from 0.0 to 1.0. The green kernel density plot represents Splam, and the blue kernel density plot represents SpliceAI-10k. **(a)** The first row shows the results of Positive-MANE, and **(b)** the second row shows the results of Positive-Alt. The first column represents the score distribution of donor sites from all splice junctions in the testing dataset, whereas the second column represents the score distribution of the acceptor sites from all splice junctions.

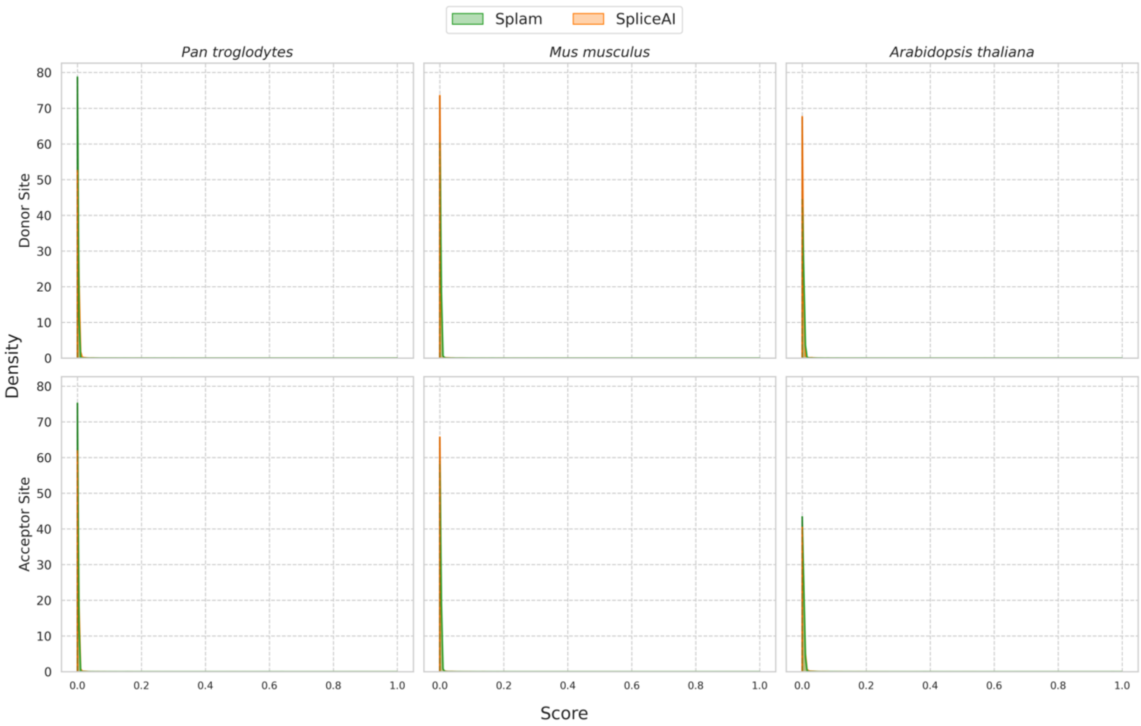


**Figure S7:** Comparison of score distributions for Splam and SpliceAI when applied to 25,000 randomly chosen splice sites from chimpanzee (left), mouse (center), and *Arabidopsis* (right). Results for donor sites are shown across the top, and acceptor sites on the bottom. Scores assigned by each program are plotted along the x-axis, while densities for both donor and acceptor sites are plotted on the y-axis. The scores for both Splam (green) and SpliceAI (orange) exhibit a narrow distribution that peaks near 0.0, indicating that the vast majority of random GT-AG splice-junction-like sequences on the reverse strand of protein-coding genes are assigned low scores, with *Arabidopsis thaliana* having a relatively lower peak.

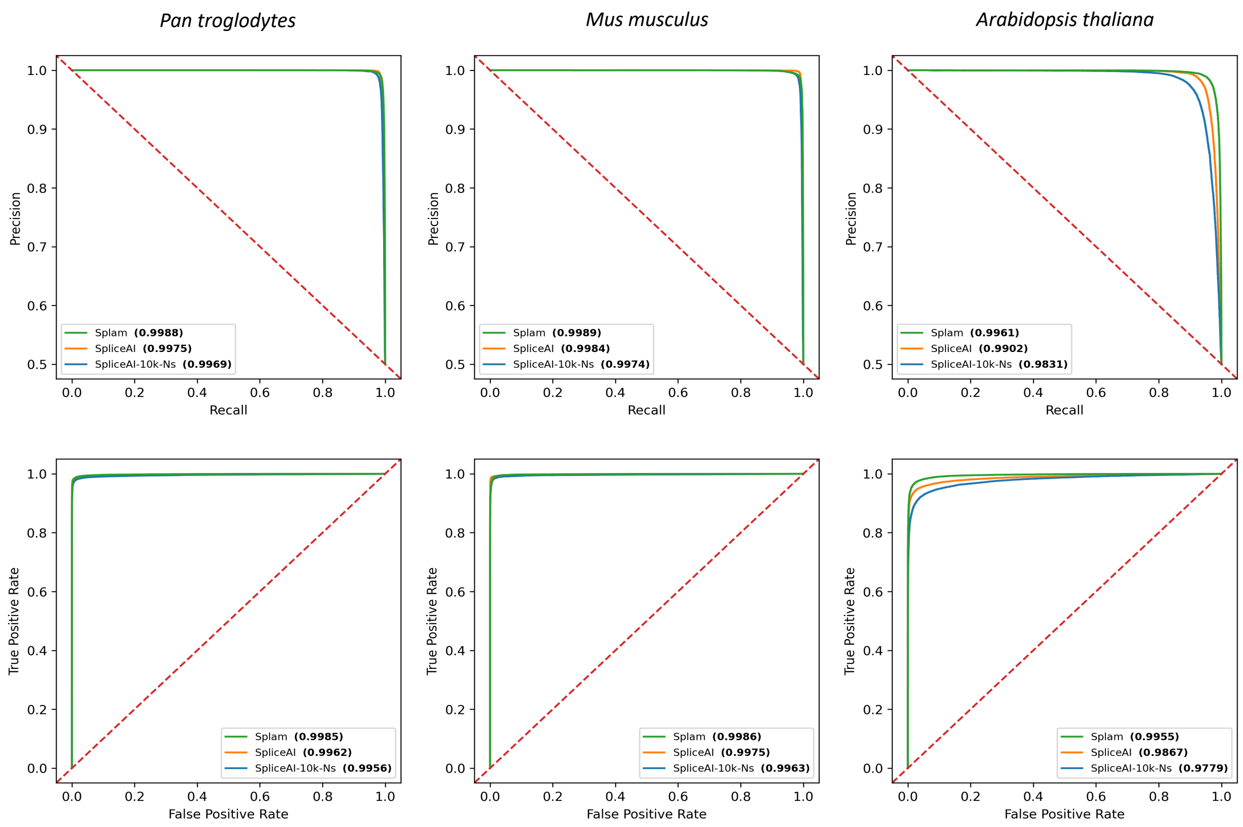


**Figure S8:** The ROC and PR curves of Splam, SpliceAI, and SpliceAI-10k-Ns were evaluated on test sets of splice junctions from three different species: chimpanzee (*Pan troglodytes*), house mouse (*Mus musculus*), and flowering plant *Arabidopsis thaliana*. Each test set consisted of 25,000 randomly selected splice junctions from its annotation file and 25,000 randomly generated pseudo-splice junctions. The pseudo-splice junctions were created by randomly selecting GT-AG pairs on the reverse strand of protein-coding genes (see Methods). The first row displays the PR curves, while the second row shows the ROC curves. We observed that Splam outperformed SpliceAI on all three datasets, and its performance was substantially better than both SpliceAI and SpliceAI-10k-Ns in the flowering plant *Arabidopsis thaliana*. For *Arabidopsis*, the AUCPR values for Splam, SpliceAI, and SpliceAI-10k-Ns are 0.996, 0.990, and 0.9831, respectively; meanwhile, the AUROC values for Splam, SpliceAI, and SpliceAI-10k-Ns are 0.996, 0.987, and 0.978, respectively.

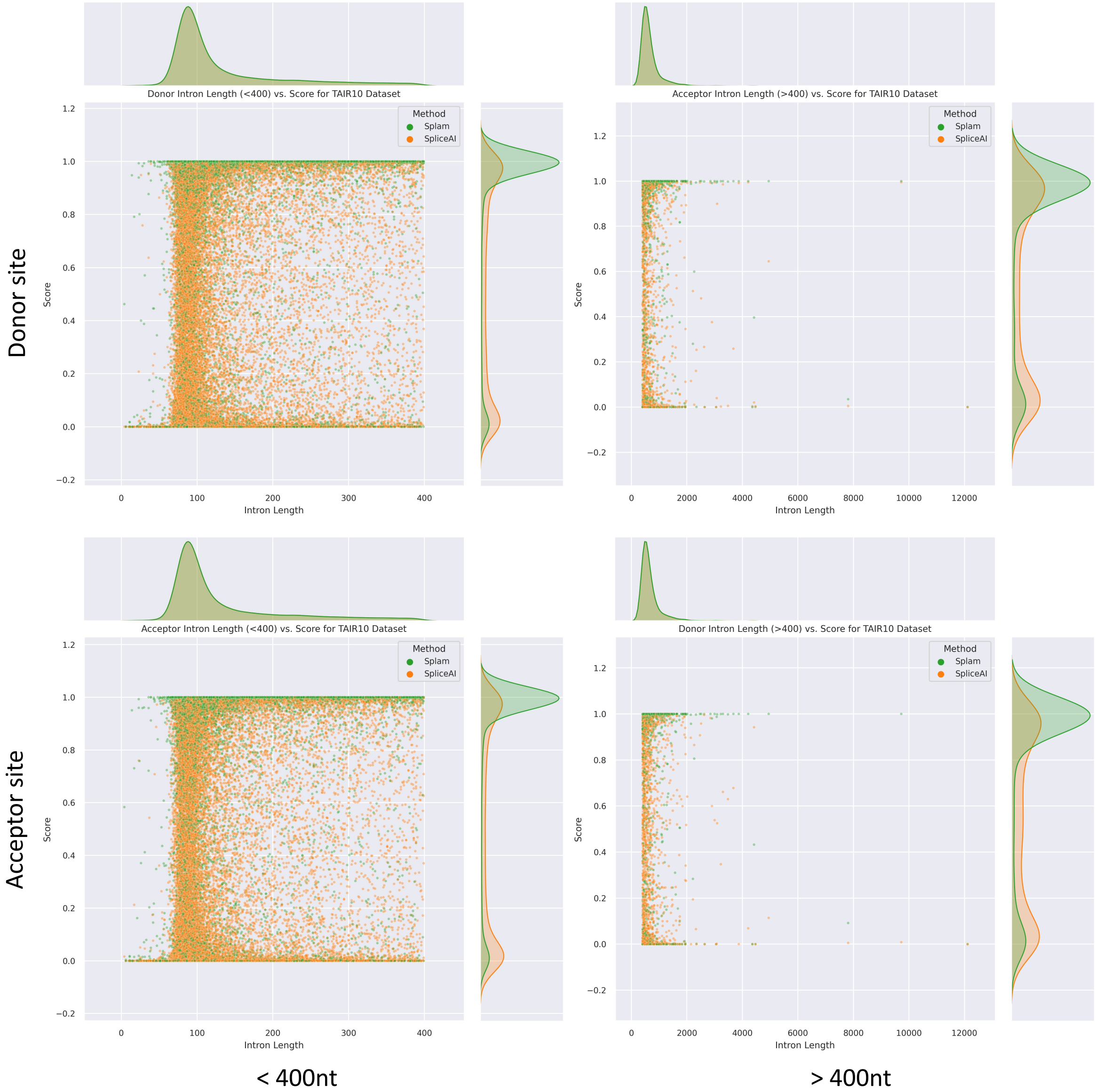


**Figure S9:** A joint scatterplot visualizing the relationship between intron lengths and score for Splam (green) and SpliceAI (orange) on the TAIR10 dataset. The left column displays intron lengths below 400nt, while the right column displays lengths above 400nt. The 400nt threshold reflects the minimum length necessary to avoid sequence overlap between the 200nt-flanked donor and acceptor sites for Splam. It additionally provides filtering of short introns for better comparison to the longer mammalian datasets. The top row shows donor site scores and the bottom row acceptor site scores. The rectangular histogram plots display the marginal distributions along each axis. The marginal score distributions do not significantly change from <400nt to >400nt for either model, indicating that TAIR10's large discrepancy between Splam and SpliceAI scores cannot be simply attributed to shorter intron lengths.

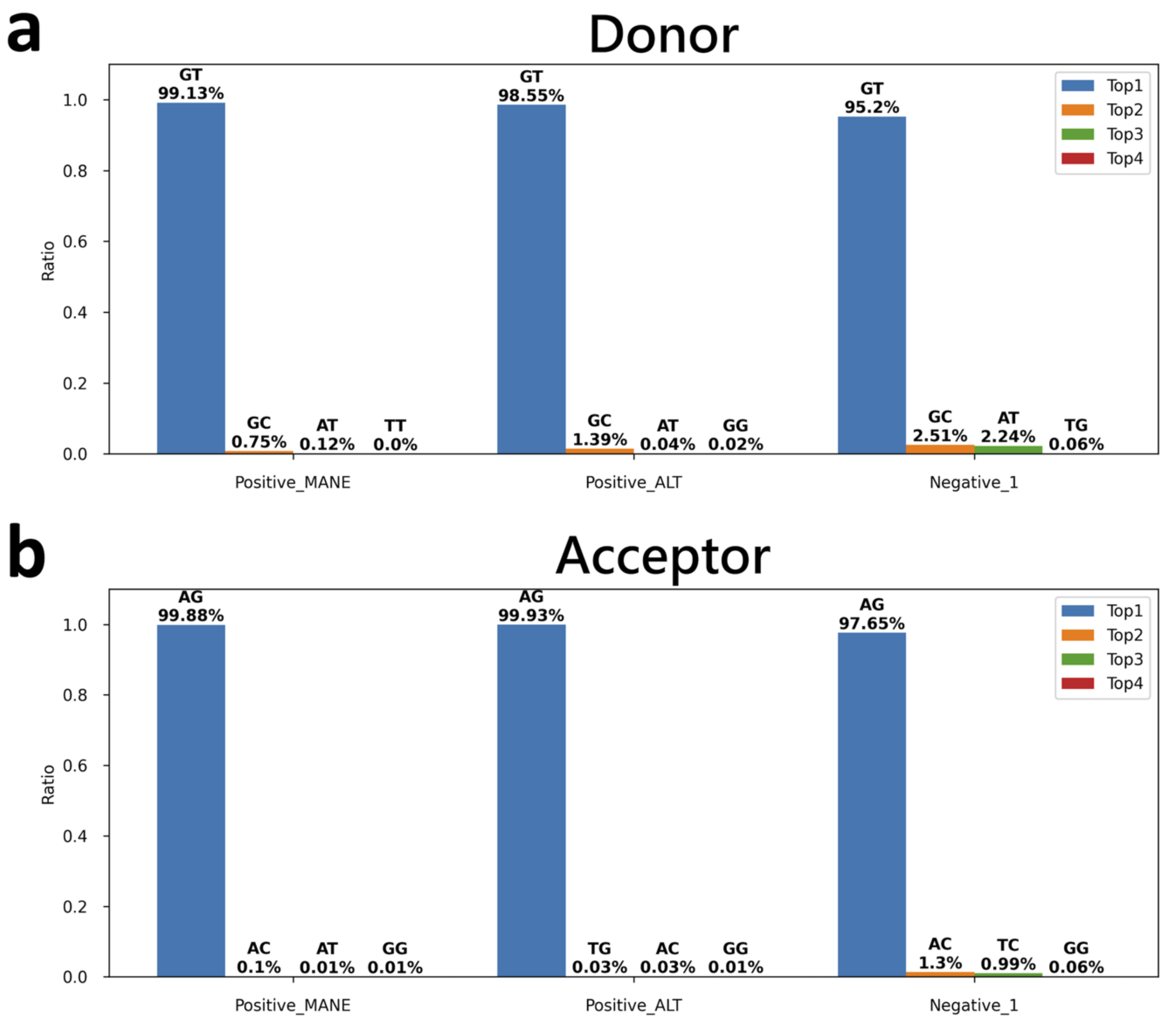


**Figure S10:** The four most frequently-occuring dinucleotides, by proportion, for the **(a)** donor and **(b)** acceptor sites for Positive-MANE, Positive-Alt, and Negative-1. It is observed that the canonical GT-AG donor-acceptor pair predominates across all three datasets with Negative-1 has the lowest frequency.

| Tools | | Accuracy (%) (Donor / Acceptor / Junction) | Recall (%) (Donor / Acceptor / Junction) | Precision (%) (Donor / Acceptor / Junction) |
|---|---|---|---|---|
| *Pan troglodytes* | Splam | 97.6 / 97.6 / 97.5 | 95.3 / 95.4 / 95.1 | 99.9 / 99.8 / 99.9 |
| | SpliceAI | 95.6 / 95.9 / 93.4 | 91.6 / 92.2 / 86.8 | 99.6 / 99.7 / 99.9 |
| *Mus musculus* | Splam | 96.9 / 96.9 / 96.7 | 94.0 / 94.0 / 93.6 | 99.8 / 99.8 / 99.8 |
| | SpliceAI | 93.5 / 94.8 / 90.6 | 87.3 / 89.9 / 81.1 | 99.7 / 99.6 / 99.9 |
| *Arabidopsis thaliana* | Splam | 95.0 / 94.9 / 94.7 | 90.3 / 90.1 / 89.6 | 99.6 / 99.6 / 99.7 |
| | SpliceAI | 87.0 / 84.4 / 78.6 | 74.3 / 69.6 / 57.3 | 99.6 / 98.9 / 99.9 |

**Table S1**: The accuracy, recall, and precision of donor sites, acceptor sites, and splice junctions at the score threshold of 0.1 for Splam and SpliceAI in chimpanzee (*Pan troglodytes*), mouse (*Mus musculus*), and the flowering plant *Arabidopsis thaliana*.

| Sample | Precision | | | Recall | | |
|---|---|---|---|---|---|---|
| | Original (%) | Cleanup (%) | Δ (%) | Original (%) | Cleanup (%) | Δ (%) |
| R2826 | 51.28 | 71.84 | 20.56 ↑ | 60.00 | 59.31 | -0.69 ↓ |
| R2835 | 56.01 | 74.51 | 18.50 ↑ | 58.80 | 58.16 | -0.64 ↓ |
| R2839 | 49.65 | 64.01 | 14.36 ↑ | 62.96 | 62.16 | -0.81 ↓ |
| R2845 | 54.78 | 74.02 | 19.24 ↑ | 58.48 | 57.85 | -0.63 ↓ |
| R2855 | 45.91 | 65.47 | 19.56 ↑ | 59.63 | 58.97 | -0.66 ↓ |
| R2857 | 51.69 | 70.86 | 19.17 ↑ | 57 29 | 56.72 | -0.57 ↓ |
| R2869 | 42.44 | 66.84 | 24.40 ↑ | 58.97 | 58.25 | -0.73 ↓ |
| R2874 | 36.68 | 61.59 | 24.91 ↑ | 61.18 | 60.40 | -0.78 ↓ |
| R2894 | 60.19 | 77.36 | 17.17 ↑ | 57.91 | 57.33 | -0.58 ↓ |
| R2895 | 50.70 | 71.73 | 21.03 ↑ | 60.23 | 59.53 | -0.70 ↓ |

**Table S2**: The precision and recall scores at the intron level for the 10 poly-A captured samples.

| Sample | Precision | | | Recall | | |
|---|---|---|---|---|---|---|
| | Original (%) | Cleanup (%) | Δ (%) | Original (%) | Cleanup (%) | Δ (%) |
| R12258 | 51.31 | 80.36 | 29.05 ↑ | 60.16 | 59.33 | -0.83 ↓ |
| R12260 | 37.92 | 74.45 | 36.53 ↑ | 63.23 | 62.16 | -1.07 ↓ |
| R12263 | 44.71 | 78.05 | 33.34 ↑ | 61.43 | 60.50 | -0.93 ↓ |
| R12265 | 45.74 | 79.75 | 34.02 ↑ | 60.03 | 59.15 | -0.88 ↓ |
| R12266 | 43.31 | 76.66 | 33.35 ↑ | 62.24 | 61 25 | -0.99 ↓ |
| R12277 | 53.07 | 85.11 | 32.04 ↑ | 57.06 | 56.33 | -0.73 ↓ |
| R12278 | 43.17 | 76.89 | 33.72 ↑ | 61 95 | 61.01 | -0.94 ↓ |
| R12280 | 40.93 | 74.99 | 34.06 ↑ | 62.77 | 61.80 | -0.98 ↓ |
| R12285 | 58.52 | 86.06 | 27.53 ↑ | 55.85 | 55.20 | -0.65 ↓ |
| R12287 | 39.36 | 73.19 | 33.83 ↑ | 63.75 | 62.68 | -1.07 ↓ |

**Table S3**: The precision and recall scores at the intron level for the 10 ribosomal RNA depletion samples.

| Sample | Precision | | | Recall | | |
|---|---|---|---|---|---|---|
| | Original (%) | Cleanup (%) | Δ (%) | Original (%) | Cleanup (%) | Δ (%) |
| R2826 | 39.2 | 41.6 | 2.4 ↑ | 10.6 | 10.6 | 0.0 |
| R2835 | 41.4 | 43.5 | 2.1 ↑ | 10.1 | 10.1 | 0.0 |
| R2839 | 37.7 | 40.3 | 2.6 ↑ | 11.1 | 11.1 | 0.0 |
| R2845 | 38.8 | 40.9 | 2.1 ↑ | 10.1 | 10.1 | 0.0 |
| R2855 | 39.4 | 41.8 | 2.4 ↑ | 10.4 | 10.4 | 0.0 |
| R2857 | 38.8 | 41.0 | 2.2 ↑ | 95 | 9.5 | 0.0 |
| R2869 | 37.9 | 40.4 | 2.5 ↑ | 10.1 | 10.1 | 0.0 |
| R2874 | 36.6 | 39 3 | 2.7 ↑ | 10.9 | 10.9 | 0.0 |
| R2894 | 40.5 | 42.8 | 2.3 ↑ | 9.9 | 9.9 | 0.0 |
| R2895 | 40.7 | 43.0 | 2.3 ↑ | 10.8 | 10.8 | 0.0 |

**Table S4**: The precision and recall scores at the transcript level for the 10 poly-A captured samples.

| Sample | Precision | | | Recall | | |
|---|---|---|---|---|---|---|
| | Original (%) | Cleanup (%) | Δ (%) | Original (%) | Cleanup (%) | Δ (%) |
| R12258 | 30.0 | 36.1 | 6.1 ↑ | 9.2 | 9.4 | 0.2 ↑ |
| R12260 | 24.7 | 31.0 | 6.3 ↑ | 9.5 | 9.9 | 0.4 ↑ |
| R12263 | 27 1 | 33.1 | 6.0 ↑ | 9.4 | 9.6 | 0.2 ↑ |
| R12265 | 25.9 | 31.9 | 6.0 ↑ | 8.7 | 8.9 | 0.2 ↑ |
| R12266 | 27.1 | 32.9 | 5.8 ↑ | 9.7 | 9.9 | 0.2 ↑ |
| R12277 | 25.9 | 31.2 | 5.3 ↑ | 7.4 | 7.6 | 0.2 ↑ |
| R12278 | 27.9 | 34 4 | 6.5 ↑ | 97 | 10.0 | 0.3 ↑ |
| R12280 | 27.0 | 33.4 | 6.4 ↑ | 9.9 | 10.2 | 0.3 ↑ |
| R12285 | 26.0 | 32 2 | 6.2 ↑ | 7.1 | 73 | 0.2 ↑ |
| R12287 | 26.0 | 31.5 | 5.5 ↑ | 9.9 | 10.2 | 0.3 ↑ |

**Table S5**: The precision and recall scores at the transcript level for the 10 ribosomal RNA depletion samples.

| Sample | Matching intron chains | | | Matching transcripts | | |
|---|---|---|---|---|---|---|
| | **Original** | **Cleanup** | **Δ (%)** | **Original** | **Cleanup** | **Δ (%)** |
| R2826 | 21,594 | 21,596 | 0.01 ↑ | 21,858 | 21,861 | 0.02 ↑ |
| R2835 | 20,697 | 20,647 | -0.24 ↓ | 20,958 | 20,915 | -0.21 ↓ |
| R2839 | 22,801 | 22,789 | -0.05 ↓ | 23,024 | 23,012 | -0.05 ↓ |
| R2845 | 20,612 | 20,602 | -0.05 ↓ | 20,875 | 20,870 | -0.02 ↓ |
| R2855 | 21,291 | 21,262 | -0.14 ↓ | 21,526 | 21,508 | -0.08 ↓ |
| R2857 | 19,393 | 19,364 | -0.15 ↓ | 19,629 | 19,596 | -0.17 ↓ |
| R2869 | 20,532 | 20,493 | -0.19 ↓ | 20,835 | 20,810 | -0.10 ↓ |
| R2874 | 22,187 | 22,180 | -0.03 ↓ | 22,478 | 22,484 | 0.03 ↑ |
| R2894 | 20,236 | 20,265 | 0.14 ↑ | 20,481 | 20,514 | 0.16 ↑ |
| R2895 | 2,157 | 22,105 | -0.23 ↓ | 22,420 | 22,369 | -0.23 ↓ |

**Table S6**: The number of matching intron chains and transcripts for the 10 poly-A captured samples.

| Sample | Matching intron chains | | | Matching transcripts | | |
|---|---|---|---|---|---|---|
| | **Original** | **Cleanup** | **Δ (%)** | **Original** | **Cleanup** | **Δ (%)** |
| R12258 | 18,869 | 19,267 | 2.1 ↑ | 19,071 | 19,500 | 2.3 ↑ |
| R12260 | 19,562 | 20,285 | 3.7 ↑ | 19,752 | 20,513 | 3.8 ↑ |
| R12263 | 19,207 | 19,661 | 2.4 ↑ | 19,419 | 19,910 | 2.5 ↑ |
| R12265 | 17,799 | 18,291 | 2.8 ↑ | 17,995 | 18,513 | 2.9 ↑ |
| R12266 | 19,838 | 20,281 | 2.3 ↑ | 20,040 | 20,525 | 2.4 ↑ |
| R12277 | 15,198 | 15,439 | 1.6 ↑ | 15,384 | 15,656 | 1.8 ↑ |
| R12278 | 19,835 | 20,359 | 2.4 ↑ | 20,063 | 20,624 | 2.8 ↑ |
| R12280 | 20,293 | 20,868 | 2.9 ↑ | 20,504 | 21,114 | 3.0 ↑ |
| R12285 | 14,615 | 14,971 | 2.4 ↑ | 14,774 | 15,147 | 2.5 ↑ |
| R12287 | 20,331 | 20,796 | 2.3 ↑ | 20,532 | 21,032 | 2.4 ↑ |

**Table S7**: The number of matching intron chains and transcripts for the 10 ribosomal RNA depletion samples.

| Sample | Hypothetical exons | | | Missed exons | | |
|---|---|---|---|---|---|---|
| | Original (%) | Cleanup (%) | Δ (%) | Original (%) | Cleanup (%) | Δ (%) |
| R2826 | 7.9 | 5.7 | -2.2 ↓ | 47.7 | 47 9 | 0.2 ↑ |
| R2835 | 6.6 | 4.7 | -1.9 ↓ | 49.0 | 49.2 | 0.2 ↑ |
| R2839 | 9.7 | 6.8 | -2.9 ↓ | 45.4 | 45.7 | 0.3 ↑ |
| R2845 | 7.5 | 5.3 | -2.2 ↓ | 48.9 | 49.1 | 0.2 ↑ |
| R2855 | 7.3 | 5.0 | -2.3 ↓ | 48.3 | 48.5 | 0.2 ↑ |
| R2857 | 6.4 | 4.3 | -2.1 ↓ | 50.1 | 50.3 | 0.2 ↑ |
| R2869 | 7.7 | 5.2 | -2.5 ↓ | 48.7 | 49.0 | 0.3 ↑ |
| R2874 | 9.4 | 6.6 | -2.8 ↓ | 46.7 | 46.9 | 0.2 ↑ |
| R2894 | 6.6 | 4.7 | -1.9 ↓ | 49 4 | 49.7 | 0.3 ↑ |
| R2895 | 7.6 | 5.5 | -2.1 ↓ | 47.8 | 48.1 | 0.3 ↑ |

**Table S8**: The percentage of hypothetical exons and missed exons for the 10 poly-A captured samples.

| Sample | Hypothetical exons | | | Missed exons | | |
|---|---|---|---|---|---|---|
| | Original (%) | Cleanup (%) | Δ (%) | Original (%) | Cleanup (%) | Δ (%) |
| R12258 | 12.9 | 6.4 | -6.5 ↓ | 47.1 | 47.4 | 0.3 ↑ |
| R12260 | 18.0 | 9.1 | -8.9 ↓ | 44.6 | 45.0 | 0.4 ↑ |
| R12263 | 15.2 | 7.5 | -7.7 ↓ | 46.1 | 46.5 | 0.4 ↑ |
| R12265 | 15.3 | 7.2 | -8.1 ↓ | 47.3 | 47 6 | 0.3 ↑ |
| R12266 | 15.9 | 8.3 | -7.6 ↓ | 45.5 | 45.9 | 0.4 ↑ |
| R12277 | 12.5 | 5.4 | -7.1 ↓ | 49.6 | 49.9 | 0.3 ↑ |
| R12278 | 15.2 | 7.6 | -7.6 ↓ | 45.8 | 46.1 | 0.3 ↑ |
| R12280 | 16.8 | 8.6 | -8.2 ↓ | 45.1 | 45.5 | 0.4 ↑ |
| R12285 | 12.5 | 5.1 | -7.4 ↓ | 50.7 | 50.9 | 0.2 ↑ |
| R12287 | 16.8 | 9.2 | -7.6 ↓ | 44.3 | 44.7 | 0.4 ↑ |

**Table S9**: The precision of hypothetical exons and missed exons for the 10 ribosomal RNA depletion samples.

| Sample | Hypothetical introns | | | Missed introns | | |
|---|---|---|---|---|---|---|
| | Original (%) | Cleanup (%) | Δ (%) | Original (%) | Cleanup (%) | Δ (%) |
| R2826 | 4.2 | 2.5 | -1.7 ↓ | 35.9 | 36.3 | 0.4 ↑ |
| R2835 | 3.5 | 2.1 | -1.4 ↓ | 37.4 | 37.6 | 0.2 ↑ |
| R2839 | 5.3 | 3.2 | -2.1 ↓ | 33.6 | 34.0 | 0.4 ↑ |
| R2845 | 3.9 | 2.3 | -1.6 ↓ | 37.4 | 37 7 | 0.3 ↑ |
| R2855 | 3.9 | 2.2 | -1.7 ↓ | 36.4 | 36.7 | 0.3 ↑ |
| R2857 | 3.3 | 1.8 | -1.5 ↓ | 38.5 | 38.8 | 0.3 ↑ |
| R2869 | 4.1 | 2.2 | -1.9 ↓ | 37 5 | 37.8 | 0.3 ↑ |
| R2874 | 5.0 | 2.9 | -2.1 ↓ | 35.0 | 35.3 | 0.3 ↑ |
| R2894 | 3.4 | 2.0 | -1.4 ↓ | 37.9 | 38.2 | 0.3 ↑ |
| R2895 | 4.0 | 2.4 | -1.6 ↓ | 35.9 | 36.2 | 0.3 ↑ |

**Table S10**: The percentage of hypothetical introns and missed intron for the 10 poly-A captured samples.

| Sample | Hypothetical introns | | | Missed introns | | |
|---|---|---|---|---|---|---|
| | Original (%) | Cleanup (%) | Δ (%) | Original (%) | Cleanup (%) | Δ (%) |
| R12258 | 7.6 | 2.9 | -4.7 ↓ | 35.7 | 36.1 | 0.4 ↑ |
| R12260 | 10 5 | 3.9 | -6.6 ↓ | 33.8 | 34.0 | 0.2 ↑ |
| R12263 | 8.9 | 3.3 | -5.6 ↓ | 35.0 | 35.4 | 0.4 ↑ |
| R12265 | 8.9 | 3.0 | -5.9 ↓ | 36.4 | 36.6 | 0.2 ↑ |
| R12266 | 9.3 | 3.6 | -5.7 ↓ | 34.3 | 34 7 | 0.4 ↑ |
| R12277 | 7.2 | 2.3 | -4.9 ↓ | 38.8 | 39.1 | 0.3 ↑ |
| R12278 | 9.0 | 3.5 | -5.5 ↓ | 34.4 | 34.8 | 0.4 ↑ |
| R12280 | 9.8 | 3.8 | -6.0 ↓ | 33.8 | 34.2 | 0.4 ↑ |
| R12285 | 7.4 | 2.1 | -5.3 ↓ | 39.9 | 40.1 | 0.2 ↑ |
| R12287 | 9.9 | 4.1 | -5.8 ↓ | 33.1 | 33.4 | 0.3 ↑ |

**Table S11**: The precision of hypothetical introns and missed introns for the 10 ribosomal RNA depletion samples.

| | | | | | | |
|---|---|---|---|---|---|---|
| **Positive_MANE** | Donor | **GT** | **GC** | **AT** | **TT** | **GA** |
| | | 178618 | 1346 | 209 | 8 | 8 |
| | Acceptor | **AG** | **AC** | **AT** | **GG** | **AA** |
| | | 178874 | 184 | 14 | 10 | 9 |
| **Positive_Alt** | Donor | **GT** | **GC** | **AT** | **GG** | **GA** |
| | | 84633 | 1195 | 36 | 17 | 13 |
| | Acceptor | **AG** | **TG** | **AC** | **GG** | **AT** |
| | | 85815 | 25 | 24 | 12 | 9 |

**Table S12**: The count of each dinucleotide for both the donor and acceptor sites in the Positive-MANE and Positive-Alt datasets.
